# CK2 inhibitor CX-4945 targets EWS-FLI1 signaling network and shows therapeutic efficacy in metastatic mouse models of Ewing Sarcoma

**DOI:** 10.1101/2025.09.24.677357

**Authors:** Muhammad Daniyal, Rajesh Rajaiah, Upendarrao Golla, Marudhu Pandiyan Shanmugam, Chloe Sholler, Jeremy Hengst, Abhinav B. Nagulapally, Hannah Valensi, Lanza Matthew, Yasin Uzun, Giselle Saulnier Sholler, Chandrika Gowda Behura

## Abstract

Ewing sarcoma (ES) is an aggressive bone tumor that primarily affects children, adolescents, and young adults. EWS-FLI1 oncogenic fusion protein is indispensable for ES tumor survival and progression. Casein kinase II (CK2) is a serine/threonine kinase that plays an essential role in apoptosis, DNA damage repair, and the cell cycle. CK2 is highly expressed in ES and associated with metastatic disease and poor 5-year overall survival. Here, we show that CK2 inhibitor CX-4945 (silmitasertib) induced K48-specific ubiquitination and subsequent proteasomal degradation of EWS-FLI1. CK2 inhibition effectively altered fusion protein abundance and disrupted the ES oncogenic signaling, specifically repressing metastasis-associated gene programs. Phenotypically, CK2-depleted ES cells showed decreased migration and invasion *in vitro*. In the metastatic ES xenograft model, CX-4945 significantly suppressed tumor growth, reduced tumor burden in the lungs, and extended overall survival. CK2 genetic depletion phenocopied CX-4945 effects both *in vitro* and *in vivo*. Molecular analysis of treated tumors confirmed robust target engagement, characterized by significant decrease in CK2 substrate phosphorylation levels. CX-4945 showed synergistic cytotoxicity with Irinotecan, a commonly used chemotherapy for the treatment of relapsed ES. Our findings establish CK2 as a novel therapeutic target in ES and provide a mechanistic rationale for combining CK2 inhibitor with chemotherapy regimens. Given the established safety profile of CX-4945, these results support clinical testing of the CK2 inhibitor fusion for treatment of metastatic ES. A Phase 1/2 trial (NCT06541262) is currently evaluating CX-4945 in combination with chemotherapy for pediatric and young adults with relapsed or refractory solid tumors, including ES.

**Graphical Abstract:** 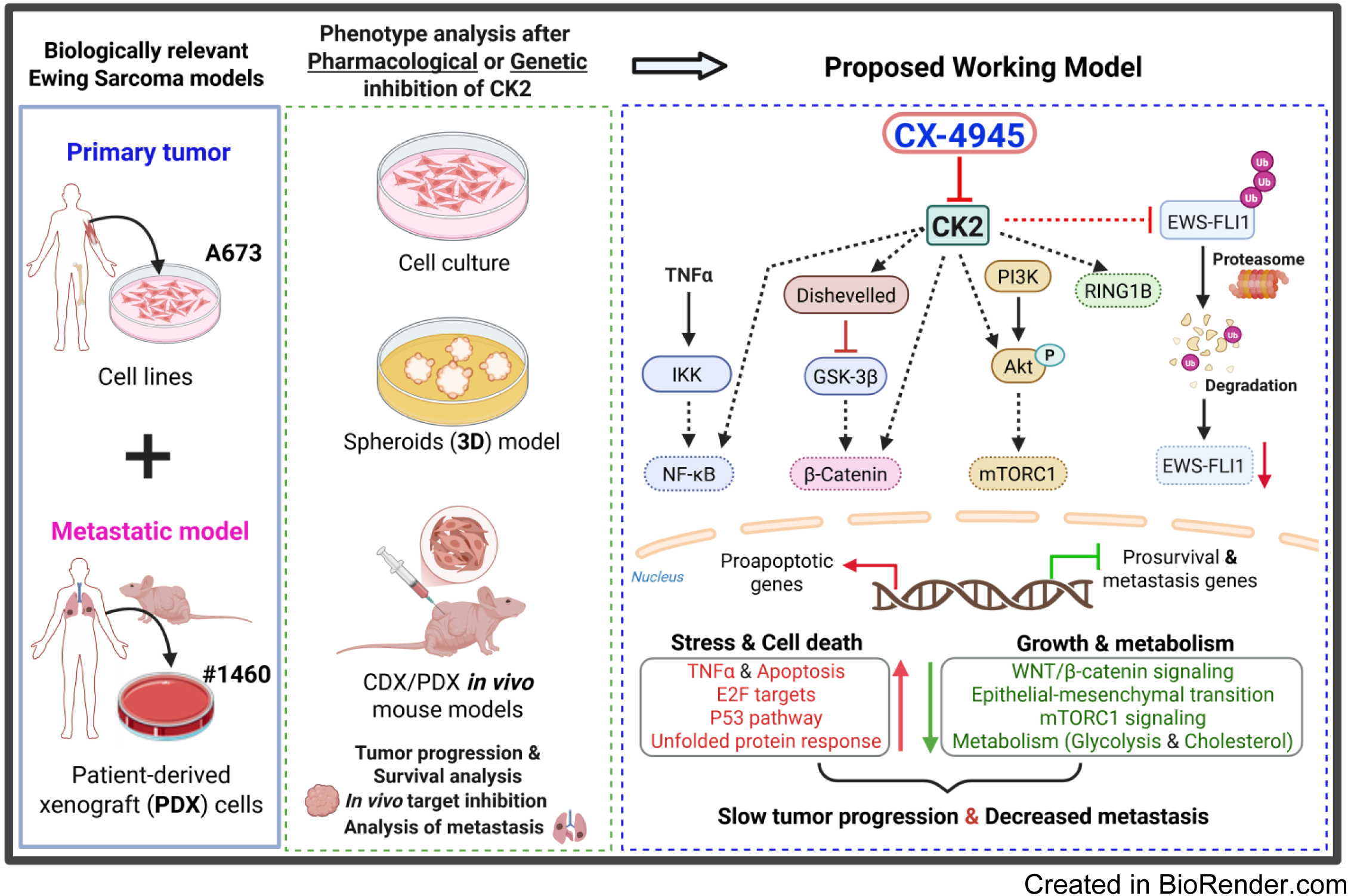

**Statement of Translational Relevance:** Our study identifies Casein Kinase 2 (CK2) as a novel therapeutic target in Ewing Sarcoma (ES). We demonstrate that CK2 inhibition triggers K48-specific ubiquitination and subsequent proteasomal degradation of EWS-FLI1 oncoprotein. Additionally, CX-4945 simultaneously targets multiple oncogenic signaling pathways and EWS-FLI1 regulators, resulting in sustained suppression of proliferation and metastasis. In metastatic ES models, oral CX-4945 showed robust efficacy, significantly reducing tumor volume and lung metastasis while extending survival. These findings provide the mechanistic rationale for integrating CK2 inhibition into current chemotherapy regimens. The translational impact is immediate: CX-4945 has an established clinical development pathway, and its safety in combination with chemotherapy is currently being evaluated in an ongoing Phase 1 multicenter trial (NCT06541262), offering a novel targeted strategy for patients with metastatic Ewing Sarcoma.

## Introduction

Ewing Sarcoma (ES) is an aggressive malignancy and the second most common bone tumor in the pediatric population (1). Despite intensive multimodal treatment, patients with metastatic or recurrent disease face a dismal prognosis, with 5-year overall survival rates remaining below 30% (2,3). While the current standard of care—comprising surgical resection, radiotherapy, and interval-compressed multi-agent chemotherapy—has improved outcomes for localized disease, there is an urgent unmet need for novel, safe, and clinically translatable therapeutic strategies for metastatic cancer (4).

Almost all ES cases harbor a pathognomonic fusion protein that drives an oncogenic transcriptional network (5,6). *EWSR1* (Ewing Sarcoma Breakpoint Region 1) is located on chromosome 22q12 and encodes an RNA-binding protein that normally helps regulate transcription and RNA processing. The *FLI1* gene, located on chromosome 11q24, encodes a member of the ETS transcription factor family. In approximately 85% of ES cases, a reciprocal translocation, t(11;22)(q24;q12), fuses the N-terminal part of *EWSR1* with the C-terminal DNA-binding domain of *FLI1* (5). While *EWSR1-FLI1* (EWS-FLI1) is the most common, *EWSR1* can also fuse with other ETS family members, such as *ERG* (about 10% of cases), *FEV*, *ETV1*, or *ETV4*. The EWS-FLI1 fusion protein is the primary driver of the ES aggressive phenotype and serves as the most significant negative predictive factor (2,3). As an aberrant transcription factor, EWS-FLI1 orchestrates widespread oncogenic gene activation or repression through extensive chromatin remodeling (7–9). Although this tumor-specific translocation represents an ideal molecular target, direct pharmacological inhibition of EWS-FLI1 has remained clinically elusive for decades (10). This “undruggable” nature of the fusion protein is due to its intrinsically disordered structure and the complex regulatory network that maintains its abundance and turnover. EWS-FLI1 has an unexpectedly high turnover rate, with a half-life estimated at 2-4 hours. Studies have shown that maintaining precise levels of EWS-FLI1 is vital for tumor survival; too much is toxic, and too little halts malignancy (11). More importantly, a small modulation in the protein abundance and activity of EWS-FLI1 can result in a significant change in tumor cell transcriptional signatures and phenotypic behaviors (12,13). EWS-FLI1 is primarily degraded by the ubiquitin-proteasome system, where E3 ubiquitin ligases mark it for destruction and deubiquitinases provide stabilization (14). Consequently, identifying and targeting the kinases that regulate EWS-FLI1 turnover is a high-priority research goal.

Casein kinase II (CK2) is a constitutively active serine/threonine kinase essential for fundamental cellular processes but frequently overexpressed in various malignancies (15–17). Typically existing as a heterotetrameric complex—comprising two catalytic subunits (alpha) and/or (alpha prime) and two regulatory beta-subunits—CK2 recognizes a distinct consensus phosphorylation sequence characterized by acidic residues (Ser/Thr-x-x-Glu/Asp) (18). As a multifunctional kinase, CK2 regulates signal transduction, transcriptional control, cell cycle progression, angiogenesis, and inflammatory responses (15,19,20). Remarkably, CK2 is responsible for phosphorylating over 20% of the human phosphoproteome, a broader substrate range than any other known kinase (21,22). This extensive reach suggests that cancer cells exhibit a state of “non-oncogene addiction” to CK2, depending more on its activity for survival and proliferation than their non-malignant counterparts (23). Key CK2 substrates implicated in oncogenesis include AKT, STAT3, MYC, and TP53 (24). Unlike many oncogenic drivers such as PI3K, RAF, or RAS, the genes encoding CK2 (primarily *CSNK2A1* and *CSNK2B*) do not typically harbor recurrent activating mutations. Instead, oncogenic signaling is driven by the ubiquitous overexpression of the wild-type enzyme, leading to increased catalytic activity across more than 25 diverse cancers, including breast, lung, prostate, colorectal carcinomas, as well as sarcomas and leukemias (25,26). This elevation in CK2 expression and activity is frequently a marker of disease progression, therapeutic resistance, and poor clinical prognosis. Given its role in sustaining the malignant phenotype through multiple non-redundant pathways, CK2 represents a highly attractive anti-cancer target.

Silmitasertib (CX-4945) is a first-in-class, orally bioavailable tetracyclic small-molecule inhibitor designed for potent, highly selective CK2 antagonism (27,28). CX-4945 functions as an ATP-competitive inhibitor of CK2 with an inhibition constant K(i) in the low nanomolar range and an excellent selectivity profile (inhibiting only 49 of 250 tested kinases) (29). Preclinical evaluations have demonstrated broad-spectrum antiproliferative activity, the direct suppression of DNA repair pathways, and the promotion of apoptosis (30,31). CX-4945 clinical development has expanded from solid tumors to viral infections and rare pediatric diseases. A favorable safety profile is documented in completed Phase 1 trials in adult tumors (32). A phase 1/2 clinical trial is currently evaluating the safety and tolerability of CX-4945 in combination with chemotherapy in pediatric and young adult patients with relapsed/refractory solid tumors, including ES (NCT06541262). In this study, we report preclinical evidence for the robust anti-tumor activity of CX-4945 in ES metastatic models and elucidate the molecular mechanisms underlying its therapeutic efficacy.

## Results

### Elevated *CSNK2A1* Expression in Ewing Sarcoma Correlates with Advanced Disease and Poor Prognosis

To determine the clinical relevance of CK2 in ES, we analyzed *CSNK2A1* (gene encoding catalytic subunit, CK2α) mRNA expression in over 300 patients utilizing the R2: Genomics Analysis and Visualization Platform (https://r2.amc.nl/). Detailed metadata for these published datasets is provided in the Supplemental Materials (**Table S1**). *CSNK2A1* levels were significantly elevated in ES tumors compared to normal bone marrow-derived mesenchymal stem cells (n=30; **Figure 1A**). Furthermore, *CSNK2A1* expression was significantly higher in patients with metastatic disease at diagnosis—a critical negative prognostic indicator—compared to those with localized tumors (**Figure 1B**). *CSNK2A1* levels also trended higher at relapse compared to initial diagnosis (**Figure 1C**). Kaplan-Meier analysis revealed that patients with high *CSNK2A1* expression had shorter overall survival than those with low expression (**Figure 1D**). Interestingly, *CSNK2B* (gene encoding the regulatory subunit, CK2β) expression was not significantly higher in tumor cells than in mesenchymal cells (**Figure S1A**), and there was no correlation with survival (**Figure S1B**). To investigate the relationship between CK2 and the known oncogenic drivers, we performed correlation analyses. *CSNK2A1* expression was positively correlated with established EWS-FLI1 target genes, including *EZH2* (Enhancer of zeste homolog 2), *GLI1* (GLI Family Zinc Finger 1), and *DAXX* (Death Domain Associated Protein) and showed a negative correlation with *IGFBP3* (Insulin-like growth factor binding protein 3) and *TGFBR2* (Transforming Growth Factor Beta Receptor 2) (**Figure 1E** and **S1C**) (13). *RNF2* (Ring Finger Protein 2), is a known regulator of EWS-FLI1 and its expression positively corelates with *CSNK2A1.* In the analysis of publicly available global CRISPR dropout screens (33), several ES cell lines (with and without EWS-FLI1 fusion) showed stronger dependency on genes encoding CK2 catalytic (*CSNK2A1*) and regulatory (*CSNK2B*) subunits for survival (**Figure 1F-1G**, **S1D**). Moreover, CK2 essentiality is comparable to the known ES dependencies such as *FLI1* and *GLI1* (**Figure 1G**, **S1D**). We measured baseline CK2 protein level and activity (substrate phosphorylation) in a series of ES cell lines and PDX samples used in this study. Both A673 cell line and the PDX 1460 cells, derived from EWS-FLI1 fusion positive relapsed ES demonstrated robust CK2α expression and heightened catalytic activity, as evidenced by the phosphorylation of the specific CK2 substrate AKT (Ser129) (**Figure 1H, S1E**). Altogether, these findings indicate that high levels and activity of protein kinase CK2 correlate with poor survival and advanced disease in ES, and strong dependency suggests CK2 as a potential therapeutic target for the treatment of ES.

**Figure 1:**
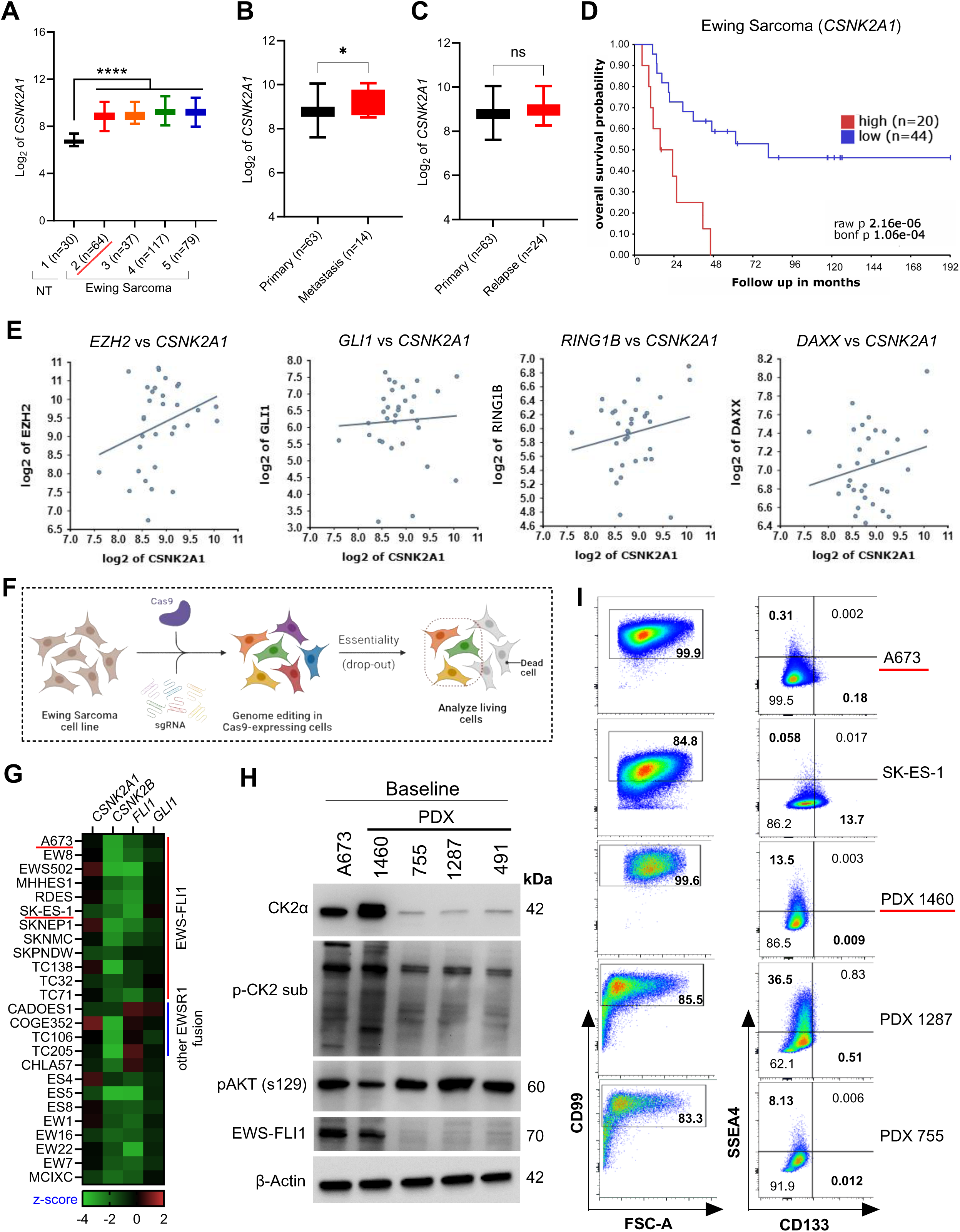
*CSNK2A1* Expression Correlates with Poor Prognosis and Advanced Disease in Ewing Sarcoma. **A)** *CSNK2A1* mRNA expression in primary Ewing sarcoma (ES) patient samples (aggregated from four datasets in R2 platform and listed in **Table S1**) compared to bone marrow mesenchymal cells -normal tissue (NT; n=30). ****p<0.0001 by mixed-effects analysis (Dunnett’s multiple comparisons test) indicates significant difference compared to NT group. **B and C)** Comparative *CSNK2A1* expression in localized versus metastatic ES (**B**) and relapse (**C**) at diagnosis. The datasets were accessed from R2 platform as listed in **Table S1**. The data are analyzed by unpaired Mann-Whitney test and *p<0.05 indicates statistical significance. ‘ns’ denotes ‘not significant’. **D)** Kaplan–Meier survival analysis of ES patients stratified by high versus low *CSNK2A1* expression. Data were analyzed using the R2: Genomics Analysis and Visualization Platform (https://r2.amc.nl/), with groups defined by median expression cutoff (n=64; GSE17679 accessed from R2 platform). Statistical significance was determined by the log-rank test (‘bonf p’ refers to the Bonferroni-corrected p-value). **E)** Gene expression correlation analysis between *CSNK2A1* and established oncogenic drivers/targets: *EZH2*, *GLI1*, *RNF2*, and *DAXX* genes in ES patients (n=64; GSE17679 accessed from R2 platform). **F)** Schematic outline of pooled CRISPR/Cas9 screens to identify genetic vulnerabilities (dropout) in ES cells. Created in BioRender.com. **G)** Heatmap summarizing the CRISPR screen z-scores (batch corrected) for indicated genes of interest in ES cell lines (n=25). Negative z-scores (dropout) indicate that targeting a specific gene caused a decrease in cell viability (i.e. essential). *FLI1* and *GLI1* genes, which are crucial for ES cells’ survival, were used as known controls. The CRISPR screening data were obtained from the iCSDB database (https://www.kobic.re.kr/icsdb/). **H)** Immunoblot analysis of baseline protein expression for CK2α, EWS-FLI1, and GLI1, alongside CK2 catalytic activity (p-AKT Ser129), in the A673 cell line and diverse ES PDX cells (1460, 755, 1287, 491). Blots from one of the representative experiments are shown. **I)** Phenotypic characterization of ES cell lines and PDX cells by flow cytometry. Baseline surface antigen profiling of three ES PDX cells and two established ES cell lines. Cells were stained with Zombie Aqua (fixable viability dye) and fluorophore-conjugated antibodies against CD99, CD133, and SSEA4. Representative dot plots show the heterogeneous expression of these markers across various models.

### Phenotypic Characterization of Ewing Sarcoma Cell Lines and PDX Models Reveals Heterogeneous Expression of Stemness and Diagnostic Markers

To establish the baseline phenotypic profiles of our experimental models, we utilized flow cytometry to quantify the surface expression of stage-specific embryonic antigen-4 (SSEA4), CD133 (prominin-1), and CD99 in two ES cell lines (A673, SK-ES-1) and three PDX models (1460, 1287, 755) (**Figure 1I**). CD99, a transmembrane glycoprotein, remains the definitive diagnostic gold standard for ES and is essential for maintaining the malignant transformed phenotype (34). SSEA4 is a glycosphingolipid marker whose high expression is functionally linked to increased proliferation, colony-forming capacity, and chemoresistance in aggressive malignancies (35). Similarly, CD133 serves as a putative marker for tumor-initiating cells (TICs) in ES, where its elevation correlates with therapeutic evasion and a heightened stemness phenotype (36). Our analysis confirmed high basal CD99 expression across all models, with A673 and PDX 1460 exhibiting near-total positivity. In the A673 cell line, SSEA4 (0.31%) and CD133 (0.18%) were minimally expressed. In contrast, SK-ES-1 cells displayed CD133-positive subpopulation (13.7%) (**Figure 1I**). The PDX models exhibited a broader spectrum of SSEA4 expression, with PDX 1287 showing the highest enrichment (36.5%), followed by PDX 1460 (13.5%) and PDX 755 (8.13%). CD133 levels remained relatively low in the PDX models, ranging from 0.012% to 0.51% (**Figure 1I**). These findings highlight the heterogeneity of ES models used here to evaluate the efficacy of CX-4945 across various malignant phenotypes. PDX 1460 was derived from a metastatic lung lesion of a pediatric relapsed ES patient. To assess molecular concordance between the PDX 1460 and the original patient tumor, we compared variance-stabilized RNA-seq expression values from the primary patient tumor with the mean of three vehicle-treated PDX replicates. Global transcriptomic comparison revealed a positive correlation (Pearson r = 0.81) between patient tumor and PDX expression profiles (**Figure S1F**). Both A673 and PDX 1460 are EWS-FLI1-positive, TP53-mutated, and intact STAG2. These results validate the use of a biologically relevant animal model of metastatic ES to evaluate the therapeutic efficacy of CX-4945 (37).

### CX-4945 Attenuates the Metastatic Potential and Induces Dose-Dependent Apoptosis in Ewing Sarcoma Cells and Tumor Organoid Models *in vitro*

To assess the therapeutic potential of CK2 inhibition, we evaluated the sensitivity of two ES cell lines (A673, SK-ES-1) and five ES PDX-derived cells to CX-4945. The molecular and clinical characteristics of the ES cell lines and PDX models used in this study are detailed in the Supplemental material (**Table S2-S3**). Dose response assays revealed IC_50_ values below 10μM across all models except SK-ES-1 cells (**Figure 2A**). To evaluate the pro-apoptotic efficacy of CX-4945, we performed real-time apoptosis monitoring by the Incucyte platform using the A673 ES cell line and the PDX 1460 cells. Treatment with increasing concentrations of CX-4945 resulted in a significant, dose-dependent elevation of green fluorescent object counts, indicating a robust induction of relative apoptosis in both the established ES cell line and the PDX model (**Figure 2B-C, S2A-B**). Furthermore, real-time apoptosis monitoring using the Incucyte platform demonstrated that 5 μM CX-4945 significantly induced programmed cell death in the majority of ES PDX lines (n=15) compared to vehicle controls (**Figure 2D**). EWS-FLI fusion-positive PDXs have a lower IC_50_ compared to fusion-negative PDX (1287). These data demonstrate that ES models exhibit a heterogeneous sensitivity to pharmacological CK2 inhibition, identifying CX-4945 as a potent inducer of apoptosis across both established and patient-derived ES cells.

**Figure 2:**
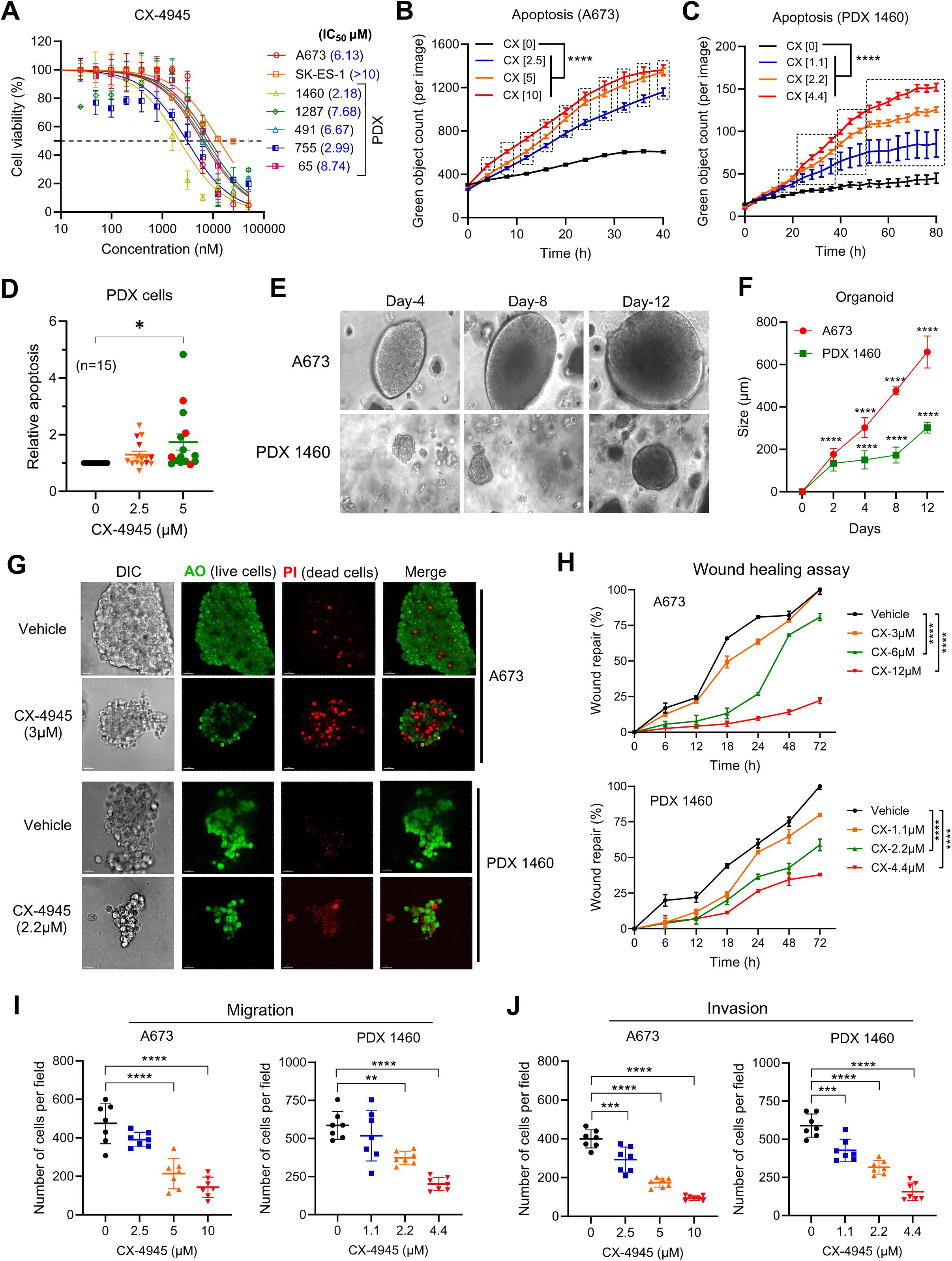
CX-4945 Induces Dose-Dependent Apoptosis and Inhibits the Metastatic Hallmarks of Ewing Sarcoma Cells. **A)** Dose-response curves illustrating the cytotoxicity of CX-4945 after 48 h treatment across ES cell lines and PDX cells, assessed via WST-1 assay. The half-maximal inhibitory concentration (IC50) of CX-4945 was determined by nonlinear regression analysis. **B and C)** Assessment of apoptosis in the A673 ES cell line (**B**) and PDX 1460 (**C**) using the Incucyte Live-Cell Analysis System. A673 (B) and PDX 1460 (C) cells were incubated in the presence or absence of CX-4945 and apoptosis was analyzed as green object count at different time points. The data are presented as mean ± SD (n=6) relative to vehicle (DMSO) control cells and analyzed by two-way ANOVA (Dunnett’s multiple comparisons test). ****p<0.0001 denotes statistical significance. **D)** Relative apoptosis of ES PDX cells (n=15) as measured by the Incucyte Live-Cell Analysis System. Data represent green fluorescent object counts at 48 h, normalized to vehicle-treated controls. *p<0.05 by one-way ANOVA (Tukey’s multiple comparisons test). **E)** Representative bright-field images of A673 and PDX 1460 organoids cultured in VitroGel 3D hydrogel system. **F)** Quantitative analysis of organoid growth/formation kinetics over a 12-day period. The data are presented as mean ± SD (n=5) and analyzed by two-way ANOVA (Sidak’s multiple comparisons test). ****p<0.0001 denotes statistical significance compared to ‘day-0’. **G)** Representative Leica SP8 confocal microscopy images of ES organoids treated with CX-4945 for 48 h. Viability was assessed using the Cyto3D Live-Dead Assay, where green fluorescence indicates live nucleated cells (Acridine orange-AO) and red fluorescence indicates apoptotic/dead cells (propidium iodide-PI). **H)** Scratch assay evaluating the migratory capacity of A673 and PDX 1460 cells. Confluent monolayers were uniformly scratched and treated with indicated concentrations of CX-4945; gap closure and repair area were monitored and recorded under brightfield microscopy at indicated time points. The data are presented as mean ± SD (n=10) relative to vehicle (DMSO) control cells. ****p<0.0001 by two-way ANOVA (Dunnett’s multiple comparisons test) indicates statistical significance. **I and J)** Anti-metastatic potential of CX-4945 was assessed by migration of ES cells (A673 and PDX 1460) in the presence or absence of CX-4945 using Transwell assay (**I**) and Invasion using Transwell assay with Matrigel (**J**) following 48 h of treatment. Data represent the mean number of cells per high-power field (HPF) averaged across 6–7 random fields. The data are presented as mean ± SD (n=7) and **p<0.01, ***p<0.001, and ****p<0.0001 by one-way ANOVA (Dunnett’s multiple comparisons test) denotes statistical significance compared to vehicle (DMSO) control treatment.

To more accurately recapitulate the complex architecture and physiological microenvironment of ES, we utilized three-dimensional (3D) tumor spheroid models derived from both the A673 cell line and PDX 1460. Tumor spheroids are increasingly recognized as superior to traditional 2D cultures for evaluating drug efficacy, as they mimic in vivo intra-tumor characteristics, such as cellular interactions and nutrient gradients (37). Following successful tumor spheroid development, growth was monitored via bright-field microscopy, with images captured every four days (**Figure 2E**). Quantitative analysis of tumor spheroid volume demonstrated consistent growth across models with PDX 1460 showing significantly slower growth kinetics compared to A673 (**Figure 2F**). To evaluate the therapeutic efficacy of CX-4945, established spheroids were treated with CX-4945 or DMSO (vehicle) for 48 hours. To assess cell viability within the 3D structure, we employed the Cyto3D Live-Dead Assay Kit and visualized live and dead cells using confocal microscope (**Figure 2G**). In both A673- and PDX-derived organoids, CX-4945 treatment induced a shift in cell viability: a significant increase in the red fluorescent signal (apoptotic/dead cells) and a concomitant reduction in the green fluorescent signal (viable cells) compared with vehicle controls (**Figure 2G**). These data substantiate the potent anti-tumor activity of CX-4945 on ES in a physiologically relevant 3D context.

The formation of distant metastases is a complex process involving cellular invasion, intravasation, migration, and eventual colonization at secondary sites (38). Given the correlation between high *CSNK2A1* expression and metastatic disease observed in patient datasets, we investigated the impact of CK2 inhibition on the metastasis of ES cells *in vitro*. Using 3D transwell assays and a 2D scratch assay, we assessed the migration and invasion capacity of ES cells in the presence and absence of CX-4945. Treatment with CX-4945 led to a significant, dose-dependent decrease in the migratory and invasive potential of A673 cells and PDX 1460 cells (**Figure 2H-J**). In addition, CX-4945 exhibited synergy in combination with Irinotecan (IRN), a standard chemotherapy agent, in A673 cells *in vitro* (**Figure S2C-E**). Collectively, these findings demonstrate that CX-4945 not only triggers apoptosis but also disrupts the functional mechanisms essential for ES metastasis.

### CK2 inhibition Promotes Transcriptional Rewiring of Key Oncogenic Signaling Pathways and EWS-FLI1 Gene Signature in Ewing Sarcoma Cells

To measure global transcriptomic changes following CK2 inhibition, we performed bulk RNA sequencing and differential gene expression analysis in ES cells treated with CX-4945. Both the ES cells (A673, PDX1460) showed significant transcriptional changes after the drug treatment, as indicated by principal component analysis (**Figure 3A**). Our analysis identified 4268 (UP-1920 genes; DOWN-2348 genes) and 7013 (UP-3231 genes; DOWN-3782 genes) genes that were differentially regulated by CX treatment at least 1.5-fold (adjusted p<0.05) in A673 and PDX1460 cells, respectively **(Figure 3B**; all genes are listed in additional **Supplementary Data**). The ES cells transcriptome showed moderate overlap in differentially expressed genes (DEGs) and indicates a heterogeneous transcriptional response after CX treatment (**Figure 3C**). Functional enrichment analyses of DEGs based on the Molecular Signatures Database (MSigDB) hallmark gene set showed that pro-apoptotic cellular stress response and DNA damage gene signatures related to TNFα signaling, p53 pathway, apoptosis, E2F targets, G2M checkpoint, and DNA damage were upregulated in CX-4945-treated cells (**Figure S3A**). Conversely, CK2 inhibition downregulated pro-survival gene signatures related to mTORC1 signaling, MYC targets, NOTCH signaling, metabolism, glycolysis, and hypoxia (**Figure S3A**).

**Figure 3:**
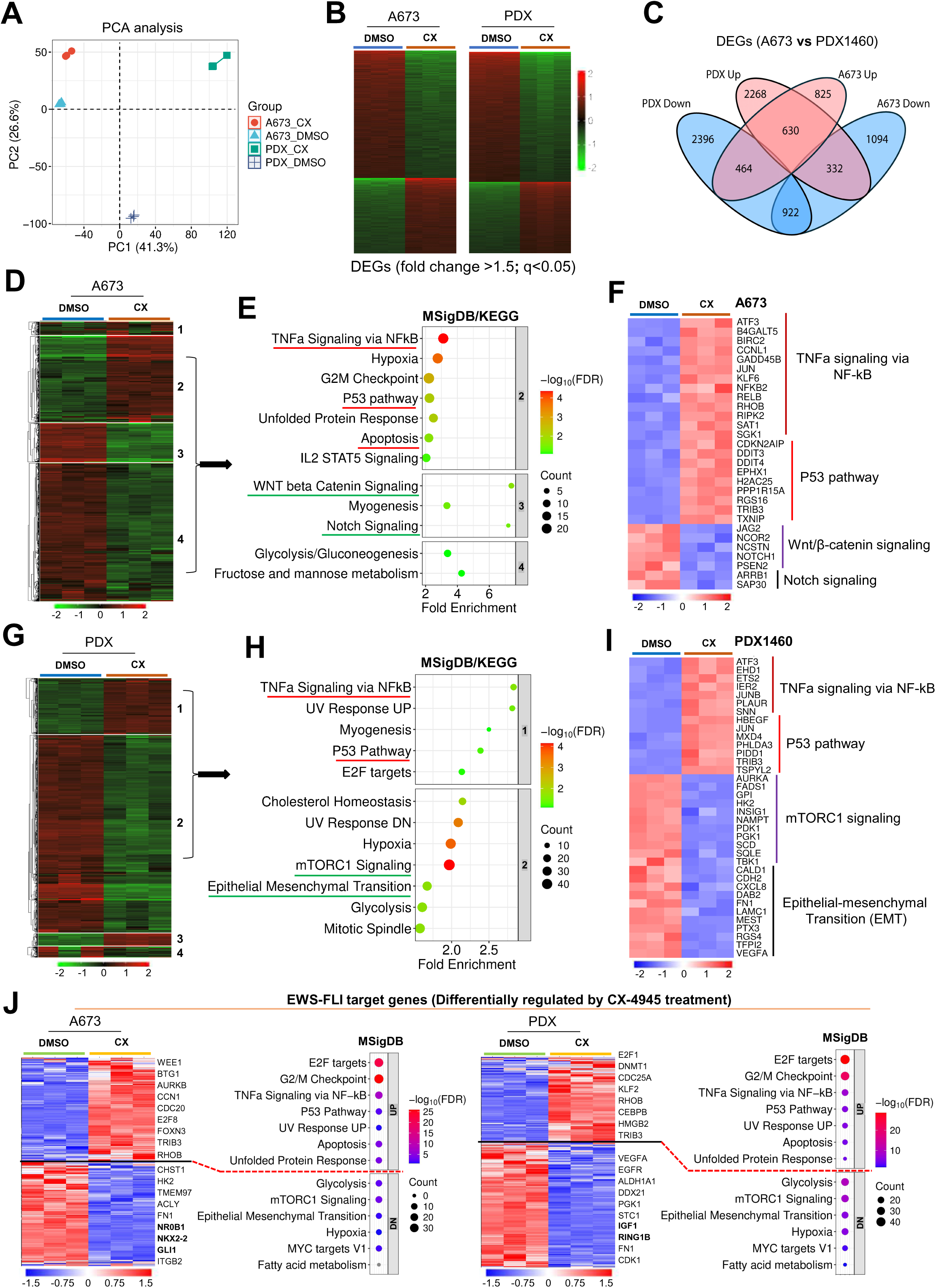
CX-4945 treatment differentially regulates cell survival pathways and EWS-FLI1 target gene signature in Ewing sarcoma cells. **A)** Principal component analysis (PCA) plot showing the distribution of ES cells (A673, PDX1460) after 24 h treatment with DMSO (vehicle) and CX-4945 (CX) (n=3). **B)** Heatmap showing the differentially expressed genes (DEGs; fold change >1.5 and adj-p <0.05) in A673 and PDX1460 ES cells following 24 h treatment with CX-4945 (n=3). **C)** Venn diagram illustrating the overlap of significant DEGs (up-and downregulated) between A673 and PDX 1460 models following CX-4945 treatment. **D)** Heatmap showing the hierarchical clustering (k-means) of the most variable genes (n=2000) in A673 ES cell line transcriptome after CX-4945 treatment for 24 h. **E)** Dot plot showing the top-ranked functional pathways (MSigDB hallmark gene set or KEGG) overrepresented in different gene clusters obtained with k-means clustering (as shown in ‘**D**’) of transcriptome in A673 cells. **F)** Heatmap showing the expression profile of few representative genes from indicated pathways enriched in different gene clusters shown in ‘**D**’. Each of the vehicle (DMSO) and CX-4945 (CX) conditions were in triplicate. **G)** Heatmap showing the hierarchical clustering (k-means) of the most variable genes (n=2000) in ES PDX1460 cells transcriptome after CX-4945 treatment. **H)** Dot plot showing the top-ranked functional pathways (MSigDB hallmark gene set or KEGG) overrepresented in different gene clusters obtained with k-means clustering (as shown in ‘**G**’) of PDX1460 transcriptome. **I)** Heatmap showing the expression profile of few representative genes from indicated pathways enriched in different gene clusters shown in ‘**G**’. Each of the vehicle (DMSO) and CX-4945 (CX) conditions were in triplicate. **J)** Heatmap showing the expression profile of EWS-FLI1 target (transcriptional mediator) genes that are differentially expressed in both A673 and PDX1460 ES cells after CX-4945 treatment. Also, the dot plot displaying the top-ranked functional pathways (MSigDB hallmark gene set) overrepresented in EWS-FLI1 target genes that were differentially regulated by CX-4945 treatment in ES cells (A673, PDX1460). ES cells showed reversal of EWS-FLI1 target gene signatures following CX-4945 treatment and substantiate potent anticancer activity of CK2 inhibitor in preclinical ES models.

Hierarchical gene cluster analysis with top variable genes in A673 transcriptome showed three distinct clusters (**Figure 3D**) that overrepresented different functional categories (**Figure 3E**). Specifically, the cluster-2 with upregulated genes is enriched for apoptosis, p53 pathway, cell cycle checkpoint, unfolded protein response, TNFα signaling, and STAT5 signaling gene signatures. The clusters 3 and 4 with genes repressed by CX treatment were enriched for WNT/ β-catenin, Notch signaling, glycolysis, and other metabolic genes (**Figure 3E and F**). Similar gene cluster analysis with the most variable genes in the PDX1460 transcriptome after CX-4945 treatment showed two distinct clusters (**Figure 3G**). The cluster-1 (genes upregulated) comprised several proapoptotic gene sets related to apoptosis, p53 pathway, E2F targets, TNFα signaling, and DNA damage (UV response UP); while the cluster-2 (repressed genes) showed enrichment of pro-survival gene signatures linked to epithelial-mesenchymal transition (EMT), mTORC1 signaling, cholesterol homeostasis, cell cycle, glycolysis, and hypoxia (**Figure 3H and I**). As both A673 and PDX1460 cells harbor the EWS-FLI1 oncoprotein, we examined relevant target gene signatures. We found that CX treatment resulted in differential regulation of several EWS-FLI1 target genes, and associated pathways indicate transcriptional rewiring of key oncogenic mediators involved in ES cells proliferation, migration, and invasion (**Figure 3J**). Specifically, EMT signature downregulation by CX-4945 treatment and CK2α genetic depletion is central to controlling ES metastatic potential and drug resistance. Taken together, our global transcriptome analysis followed by functional overrepresentation analysis revealed several important oncogenic signaling pathways and proapoptotic mechanisms targeted by CX to effectively inhibit cell proliferation and induce programmed cell death in both A673 and PDX1460 ES cells.

Next, we performed DrugSeq (Drug perturbation) enrichment analysis to compare DEGs from CX treated cells (A673 and PDX1460) against the IDG Drug Targets 2022 library. This identified several known drug-induced signatures that most closely match the transcriptomic changes caused by CX treatment **(Figure S3B).** In A673, upregulated DEGs overlapped with signatures of Multi-targeted kinase inhibitors (e.g., Midostaurin, Sorafenib) and JAK inhibitors (e.g., Ruxolitinib). This suggests that when CX inhibits CK2, the cell’s compensatory or stress response mimics the profile of these broad kinase inhibitors. Interestingly, downregulated DEGs showed enrichment for Histone Deacetylase (HDAC) inhibitor signatures (e.g., Romidepsin, Panobinostat, Belinostat), implying that CX treatment suppresses a gene expression program that is typically maintained by HDAC activity. In PDX 1460, upregulated DEGs were enriched in Purine nucleoside antimetabolite chemotherapy (e.g., Clofarabine) and Antimetabolite chemotherapy (e.g., Gemcitabine) signatures. This indicates that CX induces a stress response similar to that seen with DNA-damaging antimetabolite agents. Downregulated DEG signature overlapped with HIF-PHI (hypoxia-inducible factor prolyl hydroxylase inhibitors) and PI3Kδ inhibitors, suggesting that CX may downregulate pathways involved in hypoxia response and PI3K signaling, both of which are critical for tumor survival and progression.

### CX-4945 Reduces EWS-FLI1 Abundance via K48-Linked Proteasomal Degradation and Inhibits Downstream Oncogenic Signaling

To validate some of the molecular impacts of CK2 inhibition on the EWS-FLI1 regulatory axis seen with bulk RNA-sequencing, we measured the gene mRNA expression using quantitative RT-PCR (qRT-PCR) and protein expression by immunoblotting. Treatment with an IC50 concentration of CX-4945 led to a reduction in the mRNA expression of *EWS-FLI1* in both A673 and PDX1460 ES cells (**Figure 4A**). Furthermore, we observed significant downregulation of established EWS-FLI1 target genes (39,40), including *GLI1*, *GLI2*, *PTCH1*, *EZH2*, and *NKX2.2*, following CX-4945 treatment in both A673 cells and PDX 1460 cells (**Figure 4B, Figure S4A and B**). GLI1/GLI2/PTCH1 are the components of Hedgehog signaling pathway. EWS-FLI1 directly upregulates GLI1 which promotes tumor growth independent of canonical hedgehog ligands (41). NKX2.2 is another EWS-FLI-regulated gene that is necessary for oncogenic transformation in ES (42). EZH2 is the catalytic subunit of the Polycomb Repressive Complex 2 (PRC2) that is upregulated by EWS-FLI1 to drive epigenetic reprogramming and promote ‘stemness’ and prevent differentiation. Since EWS-FLI1 is required to keep the EZH2 promoter active, the loss of the fusion protein decreases the transcription of EZH2, allowing for potential cell differentiation or apoptosis (43). RING1B (also known as RNF2) is a critical, highly expressed protein in ES that functions as a core component of the Polycomb Repressive Complex 1 (PRC1). Correspondingly, immunoblot analysis revealed that CX-4945 dose-dependently reduced the protein levels of EWS-FLI1 and its downstream effector GLI1. This was accompanied by decreased CK2 catalytic activity—indicated by reduced p-CK2 substrate levels without altering the expression of its subunits (α/α’/β) (**Figure 4C, Figure S4C**). p-Akt (S129) is a direct CK2-specific phosphorylation site, and its reduction is a primary biomarker of CK2 activity. Beyond the direct S129 site, p-Akt (S473) also decreased significantly. This indicates loss of Akt activation, which is critical for cell survival. Dose-dependent decrease in β-Catenin levels suggests inhibition of the WNT/β-Catenin pathway, which promotes angiogenesis and tumor progression in ES. C-Myc and Cyclin D1, downstream effectors of Wnt and Akt signaling, are decreased in ES cells after CX-4945 treatment. Finally, the appearance of cleaved PARP (lower band), particularly at higher doses, is a hallmark of apoptosis (**Figure 4C**). Subcellular fractionation further confirmed that nuclear levels of GLI1 and RING1B were depleted following CK2 inhibition (**Figure S4D**).

**Figure 4:**
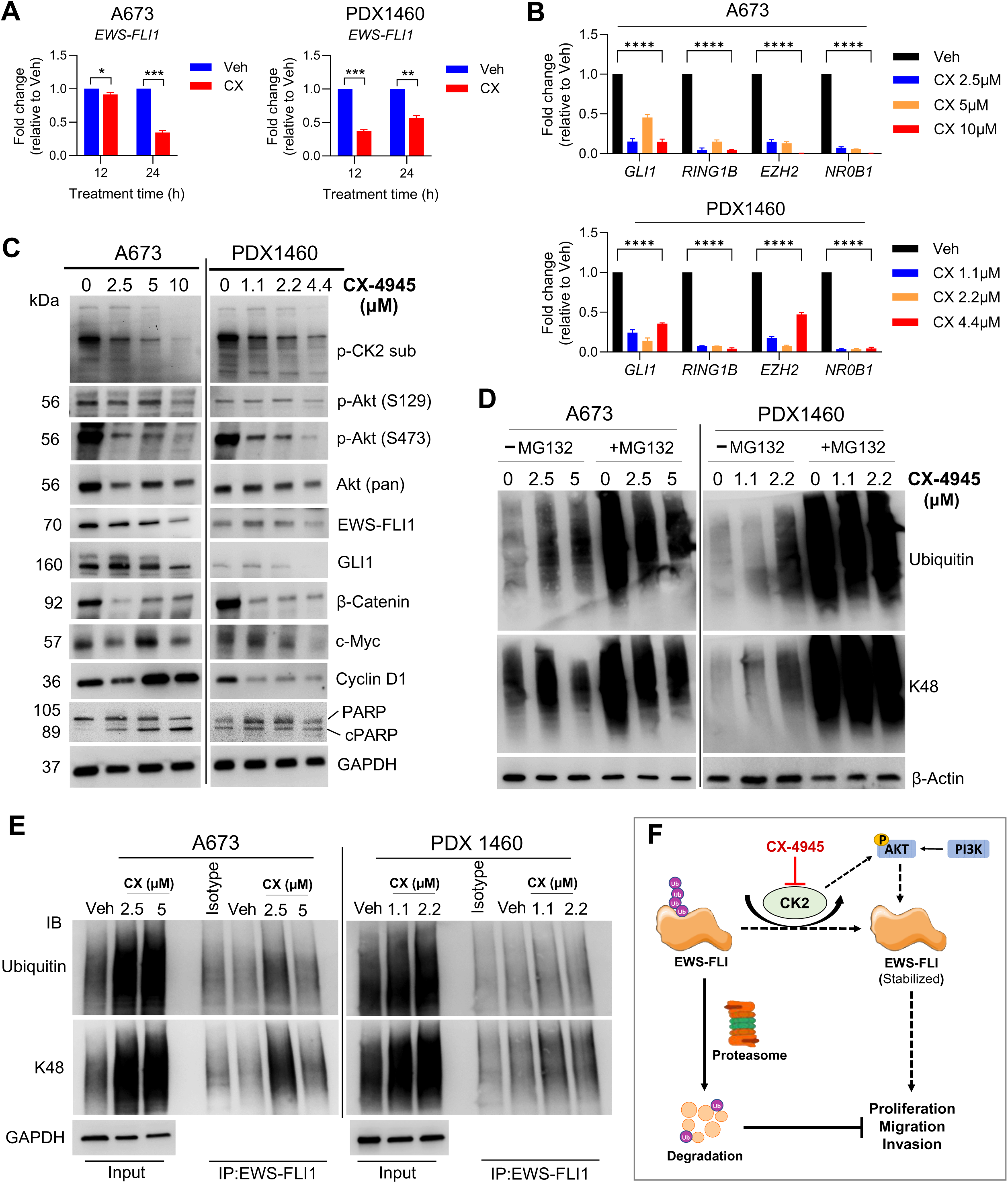
CX-4945 Downregulates the EWS-FLI1 Target Genes and Reduces Fusion Protein Abundance via K48-Linked Proteasomal Degradation. **A and B)** Quantitative RT-PCR analysis of EWSR1-FLI1 and its downstream transcriptional targets. Both A673 and PDX 1460 cells were treated with ∼IC_50_ (5µM for A673 and 2.2µM for PDX1460) dose (**A**) or indicated concentrations (**B**) of CX-4945 for 24 h and the expression of EWS-FLI1 oncogenic signature genes was quantitated by qRT-PCR. The relative gene expression was normalized to GAPDH as a housekeeping gene. The data are presented as mean ± SD (n=3 replicates from a representative run) relative to vehicle (DMSO)-treated cells. *p<0.05, **p<0.01 and ***p<0.001 by unpaired t-test (Welch’s correction) denotes statistical significance in ‘**A**’. ****p<0.0001 by two-way ANOVA (Dunnett’s multiple comparisons test) indicates statistical significance in ‘**B**’. **C)** Immunoblot analysis of EWS-FLI1 and associated signaling proteins. A673 cells (left) and PDX 1460 cells (right) were treated with increasing concentrations of CX-4945 for 24 hours. Whole-cell lysates were assessed for CK2α expression, catalytic activity (p-CK2 sub), EWS-FLI1 abundance, and downstream mediators. **D)** Western blot analysis of global and K48-linked polyubiquitination. A673 and PDX 1460 cells were treated with CX-4945 in the presence or absence of the proteasome inhibitor MG132 (10μM) for 6 h and analyzed for total and K48-linked ubiquitination level. **E**) Co-immunoprecipitation (Co-IP) of EWS-FLI1 from CX-4945-treated A673 and PDX 1460 cell lysates. EWS-FLI1 was immunoprecipitated using an anti-FLI1 antibody, and the resulting precipitates were probed for total ubiquitin and K48-specific ubiquitin to confirm targeted degradation of the fusion protein. Blots from one of the representative experiments are shown in **C-E**. **F)** Schematic diagram showing the CX-4945 proposed mechanism of action on destabilizing EWS-FLI1 fusion protein, thus inhibit proliferation, migration, and invasion capabilities of ES cells.

Given the significant reduction in EWS-FLI1 protein levels, we hypothesized that CK2 inhibition facilitates the proteasomal degradation of the fusion protein. Global protein ubiquitination analysis showed that CX-4945 treatment increased both total and K48-linked polyubiquitination in a dose-dependent manner. To confirm the degradation was proteasome-mediated, we co-treated cells with the proteasome inhibitor MG132. While MG132 effectively blocked the turnover of ubiquitinated proteins, K48-linked ubiquitin signaling remained elevated under CX-4945 treatment, suggesting that CK2 inhibition specifically marks proteins for degradation (**Figure 4D**). To establish that EWS-FLI1 is a target of ubiquitin-mediated degradation, we performed co-immunoprecipitation (Co-IP) by pulling down the EWS-FLI1 protein. Probing with ubiquitin antibodies revealed a marked increase in K48-linked polyubiquitination of EWS-FLI1 following CX-4945 treatment in both A673 and PDX1460 ES cells (**Figure 4E**). These data provide evidence that CK2 activity is essential for EWS-FLI1 stability and that CX-4945 triggers the targeted, K48-linked proteasomal degradation of this transcription factor, thereby disrupting the core oncogenic regulation of ES (**Figure 4F**).

### Genetic Inhibition of *CSNK2A1* Impairs Oncogenic Signaling, Extends Survival, and Suppresses Metastasis *In Vivo*

To validate the direct involvement of CK2 on the EWS-FLI1 regulatory axis, we performed a genetic inhibition study by transducing A673 cells with GFP-tagged lentiviral particles expressing either *CSNK2A1*-specific shRNA (shCK2α) or a scramble control (Scr). Quantitative RT-PCR (qRT-PCR) confirmed the downregulation of the EWS-FLI1-activated transcriptional program. Specifically, shCK2α cells exhibited significant transcript depletion of *EWS-FLI1*, *EZH2*, *GLI1*, *NR0B1*, *RING1B*, *NKX2.2*, *PTCH1*, and *GLI2* compared to Scr controls (**Figure 5A**, **S5A**). Immunoblot revealed reduced CK2 catalytic activity, characterized by decreased levels of p-AKT (S129), EWS-FLI1, and GLI1 protein (**Figure 5B**). Conversely, overexpression of CK2α in A673 increased CK2 substrate phosphorylation, increased expression of EWS-FLI1, and its downstream target GLI1 (**Figure 5B**). CK2α overexpression resulted in the significant upregulation of other EWS-FLI1 target genes, including GLI2, NR0B1, and NKX2.2 (**Figure S5B**). To evaluate functional differences in growth after CK2α genetic depletion, we monitored real-time proliferation using the Incucyte platform over 80 hours. CK2α knockdown (shCK2α) cells showed significantly lower proliferative indices than Scr control cells (**Figure S5C**). Furthermore, shCK2α cells displayed markedly attenuated migration, invasion, and wound-repair capacities in functional assays (**Figure 5C and D, S5D**). In a recent CRISPR screen performed by Banday et al. in A673 reporter (EWS-FLI1::TdTomato/EGFP) cells with lower EWS-FLI1 levels (tdTomato^low^ EGFP^high^), enrichment of sgRNAs targeting CK2 (*CSNK2A1*, *CSNK2A2*) aligns with our CK2 knockdown phenotype (**Figure 5E**) (44). These findings further substantiate the role of CK2 in regulating EWS-FLI1 levels in ES cells.

**Figure 5:**
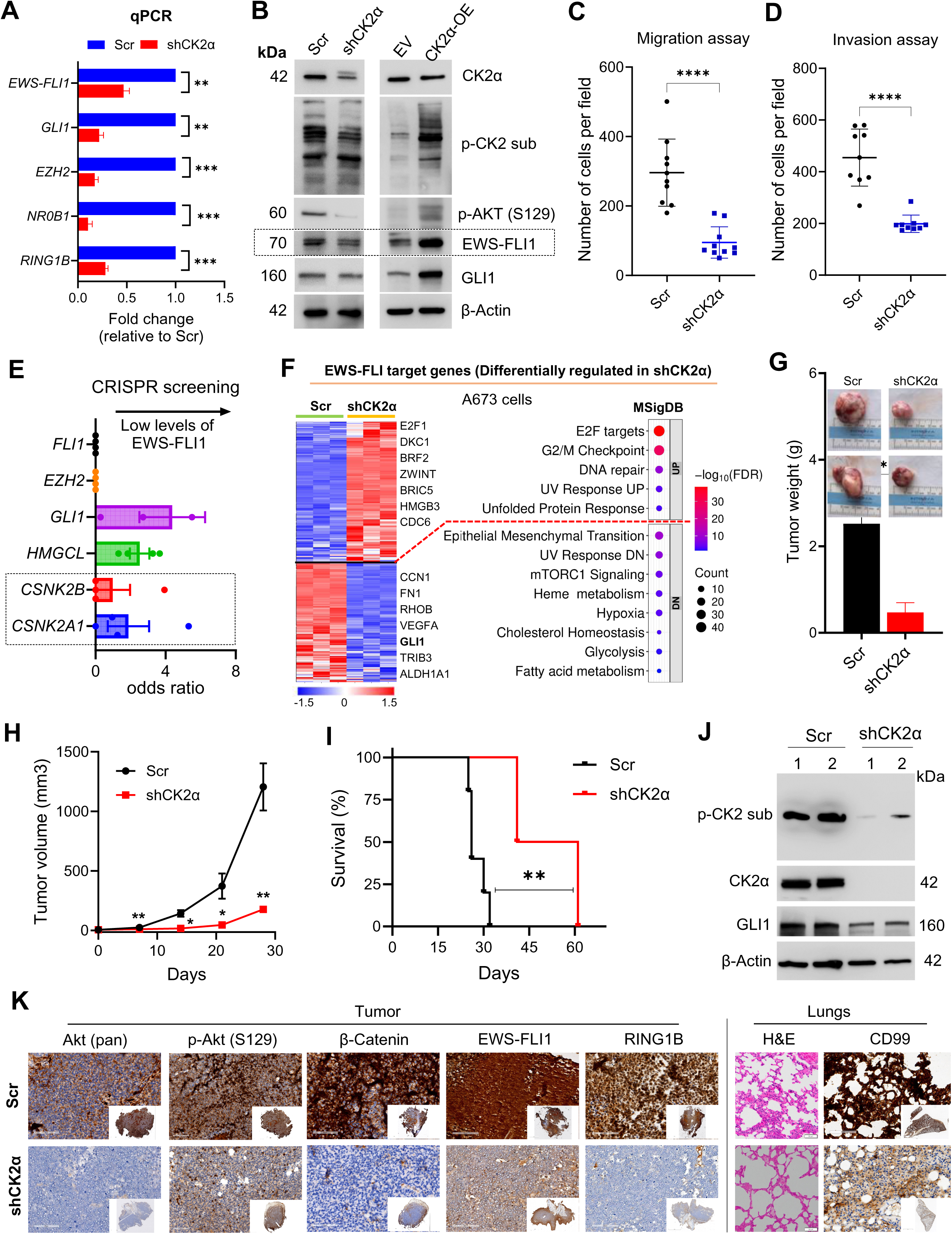
Genetic Depletion of *CSNK2A1* Impairs Oncogenic Signaling, Metastatic Potential, and Tumorigenesis in Ewing Sarcoma. **A and B)** Validation of shRNA-mediated *CSNK2A1* knockdown in A673 cells following transduction with a GFP-tagged lentiviral shRNA particles. The mRNA expression of EWS-FLI1-target genes was compared between non-targeting scramble control (Scr) and CK2α-specific shRNA (shCK2α) by qRT-PCR (**A**) and protein level via immunoblotting (**B**). The qRT-PCR data are presented as mean ± SD (n=3 replicates from a representative run) relative to Scr (scramble) control cells. **p<0.01 and ***p<0.001 by unpaired t-test (Welch’s correction) denotes statistical significance in ‘**A**’. Blots from one of the representative experiments are shown in ‘**B**’. **C and D**) Functional assessment of metastatic potential, illustrating a significant reduction in migration (**C**) and invasion (**D**) capacity following CK2α depletion. The data are presented as mean ± SD (n=9-10) and analyzed by unpaired Mann-Whitney test. ****p<0.0001 denotes statistical significance. **E**) Enrichment of sgRNAs (n=4) targeting indicated genes of interest in A673 cells sorted for EWS-FLI1 low levels relative to unsorted population in a global CRISPR screen (data adopted from Banday et al.(44)). Genes such as *EZH2*, *FLI1*, *GLI1* and *HMGCL* known to regulate EWS-FLI1 levels were used as controls. **F)** Heatmap showing the expression profile of EWS-FLI1 target (transcriptional mediator) genes that are differentially expressed in A673 ES cells after CK2α knockdown. Also, the dot plot displaying the top-ranked functional pathways (MSigDB hallmark gene set) overrepresented in EWS-FLI1 target genes that were differentially regulated by CK2α knockdown in A673 cells. ES cells showed reversal of EWS-FLI1 target gene signatures following CK2α genetic depletion and substantiate potent anticancer activity of CK2 inhibitor in ES cells. **G–I**) *In vivo* xenograft assessment of CK2α knockdown A673 cells in nude mice (n=5 per group). Comparison of terminal tumor weight quantification (inset shows a representative macroscopic image of excised tumors) (**G**), longitudinal tumor volume kinetics (**H**), and Kaplan-Meier survival analysis (**I**) of A673 xenografts. The data are presented as mean ± SEM and analyzed by unpaired t-test (Welch’s correction). *p<0.05 and **p<0.01 denotes statistical significance in ‘**G** & **H**’. **p<0.01 by Gehan-Breslow-Wilcoxon test in ‘**I**’ indicates significant difference in the overall survival of shCK2α xenograft mice. **J)** Immunoblot analysis of harvested tumor lysates evaluating CK2 catalytic activity (p-AKT Ser129) and the expression of EWS-FLI1-regulated proteins. Tumors collected from two independent mice representing each cohort are processed and presented. **K**) Histopathological evaluation of tumors by H&E staining and immunohistochemistry (IHC). Representative images of primary tumor sections (left panel) stained for CK2α, Ki67 (proliferation marker), EWS-FLI1 (anti-FLI1), and its target RNF2. Right panels illustrate H&E and CD99 staining of lung tissues between Scr and shCK2α groups. Inset image shows entire view of the tissue section.

To measure transcriptional response after CK2 genetic inhibition, we performed global transcriptome analysis in A673 cells following *CSNK2A1* knockdown (shCK2α). Our results showed 3865 genes were differentially regulated (UP-1730 genes; DOWN-2135 genes) at least 1.5-fold (adjusted p<0.05) by CK2α knockdown in A673 cells **(Figure S5E**; all genes are listed in additional **Supplementary Data**). Functional analysis of DEGs and most variable genes (cluster analysis) based on the MSigDB hallmark gene set showed that MYC targets, E2F targets, G2/M checkpoint, DNA damage, and oxidative phosphorylation were enriched in upregulated genes, whereas epithelial-mesenchymal transition (EMT), KRAS signaling, TNFα signaling via NF-κB, heme metabolism, hypoxia, and cholesterol homeostasis gene signatures were repressed in CK2α knockdown condition (**Figure S5F and G**). Moreover, genetic knockdown of CK2α altered the expression of several EWS-FLI1 target genes and associated pathways, similar to the effect seen with CX-4945 treatment (**Figure 5F**). Further, DrugSeq (drug perturbation) enrichment analysis of DEGs followed by CK2α knockdown in A673 cells revealed that transcriptional response resembled with that of other known antimetabolites (chemotherapy agents like Clofarabine and Gemcitabine), kinase inhibitors, and HDAC inhibitors (**Figure S5H**). These transcriptional responses and EWS-FLI1 gene signature after CK2α genetic knockdown corroborate with that of CK2 pharmacological blockade using CX-4945 in A673 cells.

We subsequently validated the CK2α knockdown effect *in vivo* by subcutaneously injecting Scr and shCK2α cells into athymic nude mice. Mice engrafted with shCK2α cells produced significantly smaller tumors—measured by volume and terminal weight—and achieved significantly longer overall survival than Scr controls (**Figure 5G-I**). Immunoblotting of harvested tumor cells confirmed that the *in vivo* knockdown was maintained, showing reduced CK2α abundance and diminished CK2 substrate phosphorylation (**Figure 5J**). Finally, histopathological analysis of primary tumors and lung tissues underscored the therapeutic impact of CK2 knockdown. Immunohistochemistry (IHC) of shCK2α tumors revealed reduced staining for CK2, the proliferation marker Ki67, Akt, p-Akt (S129), β-catenin, Cyclin D1, c-Myc, EWS-FLI1 and its regulator RING1B (**Figure 5K, S5I**). Notably, H&E and CD99 staining of lung sections revealed a dramatic reduction in metastatic foci in shCK2α mice compared to the extensive metastatic burden observed in Scr controls (**Figure 5K**). Collectively, these findings validate *CSNK2A1* as a critical, non-redundant driver of ES oncogenesis and metastasis.

### Pharmacological Inhibition of CK2 with CX-4945 Suppresses Tumor Progression and Systemic Metastasis *In Vivo*

The therapeutic potential of CX-4945 in a physiologically relevant context was evaluated *in vivo* using both the A673 (cell line-derived xenograft) and the PDX 1460 (patient derived xenograft) models. Athymic nude mice were subcutaneously transplanted with ES cells in the right flank and randomized to receive either CX-4945 (75 mg/kg via oral gavage) or a vehicle control (**Figure 6A**). Our results demonstrate that CX-4945 treatment resulted in significant tumor growth inhibition, leading to substantially lower terminal tumor volumes compared to the vehicle group (**Figure 6B**). Discontinuation of CX-4945 treatment of PDX1460 xenografts in a pilot study showed rapid tumor growth and thus hinted the strong anticancer potential of continued CK2 inhibition (**Figure S6A**). Terminal assessment of excised tumors further substantiated these findings, showing a marked decrease in tumor volume and tumor weight in the CX-4945 cohort (**Figure 6C and D**). Histopathological evaluation via H&E and IHC underscored the molecular impact of the treatment. Primary tumors in the CX-4945 group exhibited reduced cellularity and significantly lower staining intensity for CK2, p-AKT (S473), the proliferation marker Ki67, EWS-FLI1, and RING1B (**Figure 6E and S6B**). The reduction in primary tumor burden led to a significant increase in overall survival in both the A673 and PDX models (**Figure 6F**). Importantly, to assess the safety and translational potential of this regimen, we conducted biochemical analyses of organ function (creatinine, albumin, liver function tests) and blood count at the study endpoint. We observed no significant alterations in hematological or metabolic parameters in either the vehicle or well-tolerated without systemic toxicity (**Figure S6C**).

**Figure 6:**
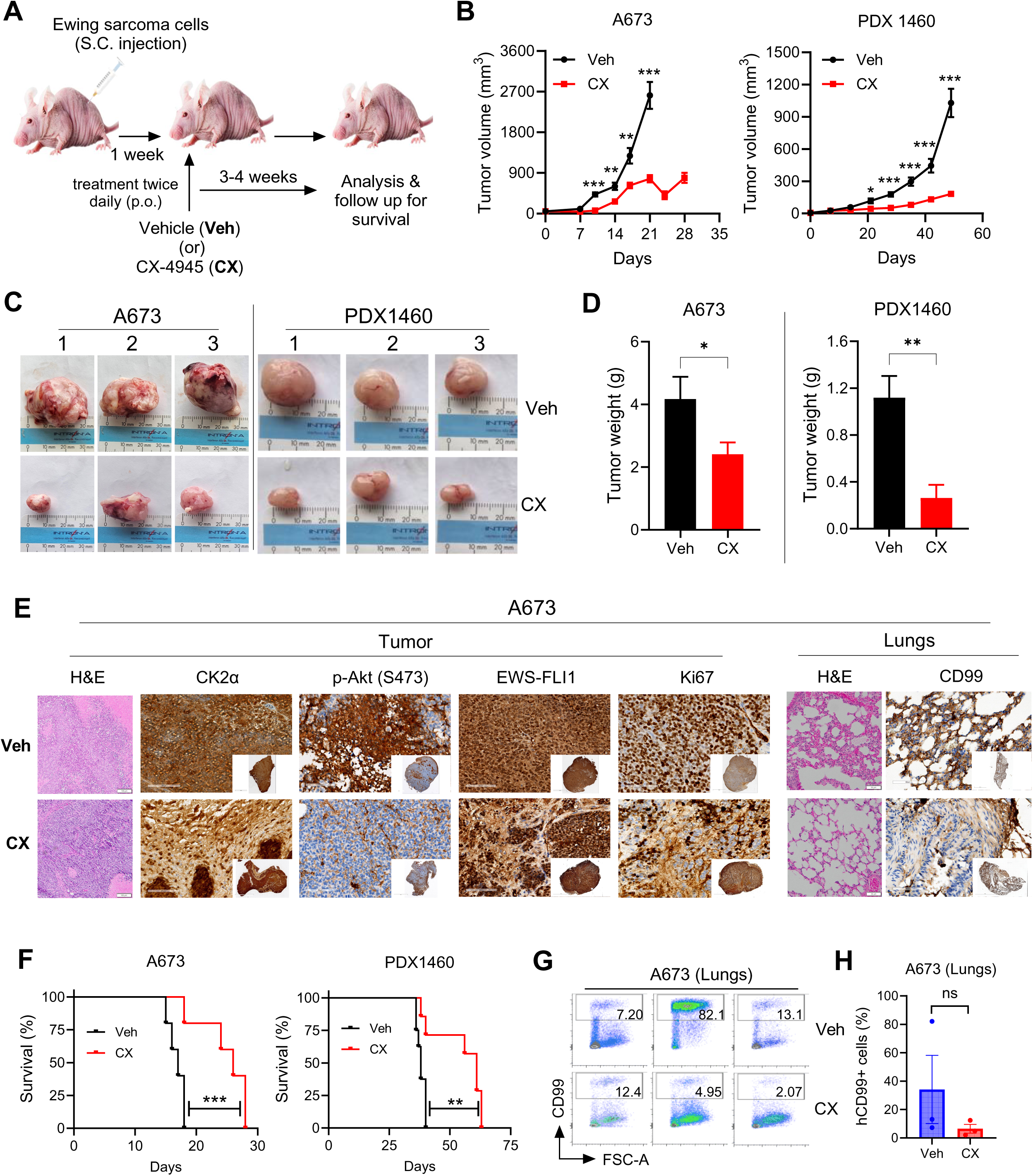
Pharmacological Inhibition of CK2 with CX-4945 Suppresses Primary Tumor Growth and Attenuates Pulmonary Metastatic Progression *In Vivo*. **A)** Schematic representation of the preclinical study design. Athymic nude mice were subcutaneously transplanted with either the A673 ES cell line or the PDX cells. Following randomization to vehicle or control group, treatment with CX-4945 (75 mg/kg BID via oral gavage) or vehicle control was started after one week. **B)** Longitudinal monitoring of tumor volume in A673 (left) and PDX 1460 (right) xenografts following Veh or CX-4945 treatment. The data are presented as mean ± SEM (n=10) and analyzed by unpaired t-test (Welch’s correction). *p<0.05, **p<0.01, and ***p<0.001 indicates statistical significance. **C and D)** Representative macroscopic images of excised tumors (**C**) and quantification of terminal tumor weight for A673 (left) and PDX 1460 (right) cohorts (**D**) with and without CX-4945 treatment. The data are presented as mean ± SEM (n=10 for A673 and n=5-7 for PDX1460) and analyzed by unpaired t-test (Welch’s correction). *p<0.05 and **p<0.01 denotes statistical significance in ‘**D**’. **E)** Histopathological and immunohistochemical (IHC) evaluation of A673 xenograft tissues. Left panels show representative primary tumor sections stained for CK2α, p-Akt (S473), EWS-FLI1 (anti-FLI1), and the proliferation marker Ki67. Right panels illustrate H&E and CD99 staining of lung tissues from Veh- and CX-4945-treated mice. Inset image shows entire view of the tissue section. **F)** Kaplan-Meier survival analysis comparing vehicle- and CX-4945-treated mice engrafted with A673 (left; n=10) and PDX 1460 (right; n=8). **p<0.01 and ***p<0.001 by Gehan-Breslow-Wilcoxon test indicates statistical significance. **G and H)** Quantitative Detection of Disseminated Tumor Cells. Flow cytometric analysis of human CD99 (hCD99)-positive cells in murine lung tissue isolated from A673 xenograft mice at the study endpoint. The data are presented as mean ± SEM (n=3). ‘ns’ denotes ‘not significant’ by unpaired t-test.

To confirm sustained target engagement at the study endpoint, we harvested primary tumors from ES-bearing mice following treatment. Immunoblot analysis of single-cell suspensions derived from excised tumors revealed a significant reduction in CK2 catalytic activity in the CX-4945-treated group, as evidenced by decreased p-CK2 substrate and CK2α levels compared to vehicle-treated controls (**Figure S6D**). However, the tumor samples were collected once the mice had reached the tumor size endpoint. We subsequently performed detailed pathological analyses of lung tissues (collected at study endpoint) to quantify the systemic metastatic burden after CX-4945 treatment. Even though both A673 and PDX 1460 represent metastatic models, the time for engraftment and time to reach endpoint were significantly longer for the PDX 1460 model. A673 mice showed significant difference in lung tumor burden between the groups while PDX 1460 had minimal lung metastasis at the study endpoint. Histopathological analysis of H&E-stained lung sections demonstrated that both the total area of metastatic foci (organized cluster of tumor cells) and the absolute number of tumor clusters (foci) were significantly higher in the vehicle control groups compared to mice receiving CX-4945 in both models (**Figure 6E, S6E-G**). The semiquantitative analysis of extra parenchymal tumor cells in the lungs showed that CX-treated mice had a lower degree of tumor cell infiltration (**Figure S6F**). 20% of CX treated mice had lung tumor foci compared to 40% in the vehicle group (**Figure S6G**). In the PDX model, one third of the vehicles showed large tumor foci, while none of the mice in the treatment group had tumor foci (**Figure S6G**). This finding is clinically significant, as metastasis remains the primary driver of mortality in ES patients. To provide a quantitative validation of pulmonary dissemination, we conducted flow cytometry analysis on mechanically dissociated lung tissue from A673 xenografts. Using the human-specific CD99 (hCD99) marker to distinguish disseminated ES cells from murine stroma, we found that vehicle-treated lungs harbored a higher frequency of hCD99+ metastatic cells than did CX-4945-treated lungs (**Figure 6G and 6H**). PDX 1460 showed minimal lung tumor burden in both groups, and the difference was not statistically significant (data not shown). Further studies incorporating local control (flank tumor resection) followed by continued follow-up and CX treatment will provide a better model for assessing the long-term effects of CX treatment on lung metastasis. Since the experiments were designed to measure tumor progression and survival difference, all mice were sacrificed when the control group mice reached the predetermined tumor endpoint. These data suggest that CK2 activity is essential not only for local tumor growth but also for the survival and distant organ colonization of circulating ES cells, identifying CX-4945 as a potent anti-metastatic agent. Collectively, these findings demonstrate that CK2 inhibition—whether via pharmacological or genetic means—suppresses metastatic progression in ES by destabilizing the core EWS-FLI1 oncogenic signaling.

## Discussion

In this study, we identified CK2 as a novel therapeutic target in Ewing Sarcoma and assessed the therapeutic efficacy of the CK2 inhibitor CX-4945 in the metastatic ES models. The single most powerful negative predictor of outcome in ES is metastatic disease (45). The metastatic process is a sequence of complex cellular events that occur in the tumor and host environments, including migration, invasion into the local microenvironment, intravasation into and extravasation from the circulation, survival of anoikis, and metastatic colonization (46). Other than EWS-FLI1 fusion, *STAG2* loss, and a few other genomic alterations, the genome is otherwise silent, suggesting non-genomic drivers and facilitators of the metastatic cascade (47). Studies have shown that fluctuations in EWS-FLI1 protein levels lead to major changes in the transcriptomic signature and cell phenotype, including metastatic potential. Cells within the same tumor transition between an actively proliferating state and a more migratory/metastatic state, and this cell plasticity is highly influenced by EWS-FLI1 turnover and activity (13). This intratumoral heterogeneity and tumor plasticity should be taken into account while determining the drug effect mediated by modulation of EWS-FLI1 dosage and direct inhibition of oncogenic signaling pathways (48). Therefore, therapies targeting EWS-FLI1 turnover and eventual phenotypic changes in ES tumors need careful consideration using biologically relevant *in vitro* and *in vivo* models. We used a primary tumor model (A673) and a metastasis model (PDX1460) to characterize the effects of CK2 inhibition on the transcriptome and cell phenotype. While both models are driven by EWS-FLI1 fusion and show aggressive metastatic phenotype, each model presents a distinct molecular signature and different cell of origin (49). CK2 inhibition significantly altered ES oncogenic programming in both models. More importantly, both *in vitro* and *in vivo* models show a significant shift in phenotype towards decreased migration, invasion, and lung metastasis.

We showed that genetic depletion of *CSNK2A1* (shCK2α) and CX-4945-mediated inhibition of catalytic CK2 subunits is sufficient to suppress pro-survival pathways and trigger apoptosis. Protein kinase CK2 operates as both a tetrameric holoenzyme (ααββ or α’α’ββ) and as independent subunits, with assembly state dictating substrate specificity. The catalytic function is redundantly managed by two isoforms, CK2α (*CSNK2A1*) and CK2α’ (*CSNK2A2*). While genetic deletion of either CK2α (*CSNK2A1*) or the regulatory subunit CK2β (*CSNK2B*) is embryonic lethal in mice, the *CSNK2A2* isoform is non-essential for viability. While both catalytic and regulatory subunits are important, CRISPR/Cas9 screens (**Figure 1**) indicate that ES cell lines often exhibit a higher dependency score for *CSNK2B* than for *CSNK2A1*. This heightened vulnerability likely stems from two factors: the functional redundancy between catalytic isoforms (where α’ may compensate for α loss) and the additional roles of CK2β. Beyond its scaffolding role in the holoenzyme, CK2β functions as a versatile regulator of alternative signaling pathways independent of CK2 catalytic activity (50). Although *CSNK2B* is overexpressed in ES tumors, its levels do not directly correlate with overall survival (**Figure S1B**), suggesting that it plays an essential role in cellular function rather than serving as a prognostic biomarker. Pharmacological targeting with CX-4945 effectively bypasses this complex subunit dynamics. CX-4945 binds the ATP-pockets of both α and α’ isoforms with high affinity, regardless of whether they are free or complexed with CK2β. By inhibiting the α subunit, CX-4945 effectively shuts down the PI3K/Akt/mTOR pro-survival axis, which is often “addicted” to CK2 activity in cancer cells. CX-4945 suppresses oncogenic signaling (e.g., Akt-S129, WNT/β-catenin) despite the continued presence of CK2β. Consequently, while ES cells show a genetic preference for CK2β loss, the catalytic inhibition provided by CX-4945 is sufficient to collapse the CK2-mediated survival network.

Our study shows that the therapeutic efficacy of CX-4945 in ES is mainly mediated by on-target CK2-mediated inhibition of the PI3K/Akt-EWS-FLI1 axis. We note that CX-4945 is a potent inhibitor of CK2, but its selectivity is notably broad, targeting several other kinases in the low-nanomolar range (51,52). Among them are DYRK1A (Dual-specificity tyrosine phosphorylation-regulated kinase 1A, regulator of cell cycle), CLK2 (CDC-like kinase 2 involved in mRNA splicing), CDK1(Cyclin-dependent kinase 1), and GSK3β (Glycogen synthase kinase 3). GSK3β is a key negative regulator of the Wnt/β-catenin pathway. DYRK1A is a master regulator of the cell cycle. CDK1 (Cyclin-Dependent Kinase 1) is a critical regulator of the cell cycle in ES, and a known therapeutic target. However, this specific off-target profile of CX-4945 can be viewed as a therapeutic advantage in ES, where multi-kinase inhibition can prevent bypass signaling and drug resistance. The simultaneous inhibition of CK2 (EWS-FLI1 stability), CLKs (splicing integrity), and PIM1 (survival signaling) creates a synergistic lethality that contributes to the therapeutic efficacy of CX-4945 in ES preclinical models. More importantly, despite its off-target reach, it is generally well-tolerated in humans, with side effects that are often manageable. The most frequent Treatment-Emergent Adverse Events (TEAEs) noted in adult cancer trials include gastrointestinal issues like diarrhea (65%), nausea (51%), and vomiting (31%) (53).

We showed that CX-4945 essentially reversed the malignant programming driven by EWS-FLI1. Treatment with the CK2 inhibitor CX-4945 elicits a significant stress-response molecular signature driving apoptosis (TNFα, UPS, E2F targets) in both models. CX-4945 treatment primarily activates the TNFα, apoptosis, and p53 pathways, likely by disrupting the oncogenic EWS-FLI1 fusion protein, which normally suppresses these tumor-suppressive signals. Additionally, the strong downregulation of hypoxia, EMT, Wnt/β-catenin, and mTORC1 signatures indicates a transition from an aggressive, metastatic state toward a therapy-responsive phenotype. ES cells use hypoxia to stabilize the EWS-FLI1 signature, driving tumor growth in low-oxygen environments (54). CX-4945 is disrupting this hypoxic adaptation, potentially re-sensitizing the ES cells to cytotoxic chemotherapy. Wnt/β-Catenin pathway promotes angiogenesis and metastasis. Consistent downregulation of Wnt/β-catenin signaling across both ES models supports the inhibition of metastasis observed in the xenograft models. Direct targets of EWS-FLI1 (e.g., *GLI1*, *EZH2*, *NR0B1*, *NKX2.2*) show a strong decrease in expression, effectively dismantling the core metastatic program of ES. RING1B is a critical modulator of EWSR1-FLI1–induced chromatin remodeling, and its inhibition is a potential therapeutic strategy for the treatment of ES (55). Proteomics analysis of Ring1B interactors has physically identified CK2 subunits as direct interactors of the RING1B complex (56). Both genetic and pharmacological inhibition of CK2 significantly decreased RING1B across all models. Our results suggest that CK2 acts as a functional partner to RING1B, regulating its oncogenic activity and facilitating the recruitment of the EWS-FLI1 complex to active enhancers. CX-4945 reduces RING1B expression and destabilizes the oncogenic EWS-FLI1 complex.

Using DrugSeq enrichment analysis, we identified a possible mechanism that resembles that of other known anti-cancer drugs. CX-4945 creates a transcriptomic state that highly resembles the effects of HDAC inhibitors (in A673) and Antimetabolites (in PDX). The overlap with multi-targeted kinase inhibitors reinforces the view that CK2 is a “master regulator” whose inhibition has broad effects on the cellular kinome and signaling landscape. The strong overlap with HDAC inhibitor signatures is particularly noteworthy as it suggests that CX-4945 might disrupt the epigenetic regulation of ES, providing a rationale for combination therapies involving CK2 and HDAC inhibitors.

We identify two possible mechanisms by which CX-4945 modulates EWS-FLI1 dosage. First, it is the CK2-mediated proteasomal degradation of EWS-FLI1. Our analysis of the K48-linkage specific ubiquitination marker revealed that CX-4945 exposure triggers a marked increase in the polyubiquitination of EWS-FLI1 (**Figure 4**). This suggests that high CK2 activity shields EWS-FLI1 from E3-ligase recognition, and when CK2 is inhibited, the fusion protein is marked for K48-linked degradation. Second, CK2-mediated inhibition of the PI3K/Akt pathway that regulates EWS-FLI1 transcription. Studies have shown that the PI3K/Akt signaling pathway is a critical upstream regulator of EWS-FLI1 transcription via the transcription factor SP1 (57). CK2 directly phosphorylates Akt at Ser129, a site that promotes Akt’s maximal activity. Ser129 is a unique, non-canonical phosphorylation site on Akt that is regulated exclusively by CK2 (58). Constitutively active CK2 maintains high Akt basal activity by phosphorylating this site. We have shown that CK2 inhibition significantly decreases p-Akt (ser129), thereby indirectly dampening the PI3K/Akt pathway and potentially reducing both the synthesis and stability of EWS-FLI1. This was shown by decreased mRNA level of EWS-FLI1 following CK2 knockdown and CX-4945 treatment.

CX-4945 induces a strong cellular stress response and drives apoptosis. The phenotypic consequences of CX-4945 treatment are a cumulative effect of EWS-FLI1 dosage-dependent and independent signaling changes. *In vitro*, shCK2-treated cells exhibited significantly slower proliferation and enhanced sensitivity to IRN-induced apoptosis. Most importantly, CK2 inhibition—both genetic and pharmacological—attenuated transwell migration and invasion potential. These findings translated to our metastatic xenograft models, where CX-4945 monotherapy significantly delayed tumor progression and extended median overall survival. Pathological evaluation of lung sections revealed that CK2 inhibition prevents the formation of micrometastatic islands. This anti-metastatic effect was quantitatively confirmed via CD99 immunohistochemistry, the clinical gold standard for identifying disseminated ES cells. In ES, where pulmonary metastasis is the primary driver of mortality, CX-4945’s ability to suppress systemic dissemination while maintaining a favorable safety profile offers a significant translational advantage. Furthermore, CX showed synergistic cytotoxicity with IRN, a commonly used cytotoxic agent in advanced ES. Given the established clinical development pathway of CX-4945, and its safety in combination with chemotherapy, is currently being evaluated in an ongoing Phase 1 multicenter trial (NCT06541262) in combination with IRN and Temozolomide.

In summary, this study identifies CK2 as a novel therapeutic target in ES, demonstrating that its inhibition triggers the proteasomal degradation of the EWS-FLI1 oncoprotein. In metastatic models, the orally bioavailable selective CK2 inhibitor CX-4945 significantly reduced tumor volume and lung metastasis while improving survival. These findings provide the mechanistic rationale for an ongoing Phase 1 multicenter trial (NCT06541262), offering a ready-to-implement targeted novel combination strategy for patients with advanced ES.

## Materials and Methods

### Cell culture

The human ES cell lines A673 (#CRL-1598, RRID:CVCL_0080) and SK-ES-1 (#HTB-86, RRID:CVCL_0627) were obtained from the American Type Culture Collection (ATCC) and cultured in Dulbecco’s Modified Eagle’s Medium (DMEM) (Corning, #10-013) for A673, McCoy’s 5A Medium (ATCC, #30-2007) for SK-ES-1, and Minimum Essential Medium α (MEMα; Gibco, #12571-063) for PDXs, growth medium supplemented with 10% heat-inactivated fetal bovine serum (FBS) (#S11150; GeminiBio, West Sacramento, CA, USA) and 1X Penicillin-Streptomycin (#15140122; Gibco, Gaithersburg, MD) at 37°C in a humidified incubator with 5% CO2. The characteristics of ES cell lines and patient-derived xenograft (PDX) cells were provided in supplementary **Table S2** and **Table S3**, respectively. Cell lines were routinely authenticated by STR profiling (Promega, Madison, Wisconsin, USA) and monitored for mycoplasma contamination. Each frozen aliquot was used and propagated for no more than 30 passages.

### Drugs and Reagents

The CK2 inhibitor CX-4945 sodium salt was generously provided by Senhwa Biosciences.

### Cell viability assays

The ES cell lines or PDX ES cells were seeded in 96-well flat-bottom plates at 1x10^5^ cells/mL in 0.1 mL/well and treated with increase concentration of CX-4945 (0.02 to 50µM) for 48 h. For drug synergy experiment, A673 cells were seeded at 1x10^5^ cells/mL in 0.1 mL/well in a 96-well flat-bottom plate and treated with increasing concentrations of Irinotecan (IRN; 0.08-5 µM) and CX-4945 (2.5-5 µM) in combination for 48 h. Following the treatment, Cell Proliferation Reagent WST-1 solution (#05015944001; Sigma-Aldrich, St. Louis, MO) was added (1:10 final dilution) and incubated for 3-4 h. Absorbance was measured at 440 nm and 650 nm (reference wavelength) by SpectraMax i3x Multi-Mode Microplate Reader (RRID:SCR_026346; Molecular Devices, San Jose, CA). Following background subtraction, absorbances were presented as % viability relative to vehicle (DMSO) control. The synergy between CX and IRN was analyzed by zero interaction potency (ZIP) reference model using the SynergyFinder Plus (https://synergyfinder.org/, RRID:SCR_019318) interactive webtool (59).

### Wound repair assay

Cells were seeded into 6-well plate and waited for the 90 to 100% confluency. After that, a uniform wound was created across the monolayer of cells using 200µL sterile pipette tip in all the conditions. All the wells were washed with PBS to remove the detached cells. Media was replaced with containing vehicle (DMSO) and CX-4945 with difference concentration for treatment groups. Closure of wound was measured over the time e.g., 0, 6, 12, 24, 48, 72 hours using microscope. Captured images were analyses for wound closure distance as % using the Image J software (RRID:SCR_003070).

### Migration assay

ES cells were serum starved for 18 h. Subsequently, 5X10^4^ cells in 500 ml of serum-free MEM alpha media were added to the upper chamber of each insert (Transwell Permeable 6.5mm insert, 8.0µM pore size, TC-treated) in a 24 well plate. 600 µL of complete media (MEM alpha with 10% FBS) were added to the lower chambers. Cell invasion chambers were incubated at 37°C, 5% CO_2_ for up to 48 h. Then, the media in the upper chamber was aspirated and any remaining media was gently removed with a cotton bud, taking care not to damage the bottom membrane of the Transwell insert. Migrated cells were fixed with 0.5% paraformaldehyde in PBS for 30 min and subsequently stained with 0.1% crystal violet dye for 10 mins, followed by washing with water for 3 to 5 times prior to air dying. Cells were viewed under light microscope, then 5 to 8 microscopic fields were imaged and counted using Image J software (RRID:SCR_003070).

### Invasion assay

Invasion assays were performed in a similar manner as the migration assays. However, Transwell inserts were coated with Corning Matrigel Basement Membrane Matrix (Cat #356234) prior to adding ES cells. The Matrigel was diluted with serum media in 1:1 ratio. 80µL of diluted Matrigel matrix coating solution was added to each Transwell insert (Corning, Cat #3422) and incubated at 37°C for two hours to facilitate polymerization. ES cells were incubated for 48 hours prior to staining. The remaining step of the protocol was identical to that of the migration assays.

### Incucyte cell proliferation and Apoptosis assays

ES cells were seeded at 5000 cells/well in 96 well plates for proliferation assay. Live-cell phase contrast images were obtained using a 10x objective lens (five images per well), and cell confluence was analyzed using IncuCyte Live Cell Analysis (v2019B) software (RRID:SCR_023147). For apoptosis assay, ES cell lines or PDX of 10,000 cells/well (50uL/well) were seeded in 96-well plates, and after 12–14 hours, Caspase 3/7 Dye for Apoptosis Green (Sartorius, Cat #440; 50uL/well with media and drug) was added. Images were obtained every 2 hours for 120 hours, and relative apoptosis was quantified using the Basic Analyzer module in Incucyte Analysis software (Sartorius 9600-0012). For apoptosis analysis in Scramble and shCK2α cells, Caspase 3/7 Dye for Apoptosis Red (Sartorius, Cat #4704) was used.

### RT-qPCR

Total RNA was isolated using TRIzol (Thermo Fisher Scientific, Waltham, MA, USA) and reverse -transcribed into cDNA using iScript cDNA Synthesis Kit (Bio-Rad Laboratories Inc., Richmond, CA). cDNA amplification was performed with PowerTrack SYBR Green Master Mix (Applied Biosystems, Vilnius, Lithuania). GAPDH was used as an internal control gene for normalization and relative expression levels were determined by the 2−ΔΔCt method [39]. The thermal cycling program consisted of the following steps: denaturation at 94 °C for 30 s and annealing/extension for 30 s at 60 °C for a total of 40 cycles. The sequences for primers used in this study were listed in **Table S4**.

### Co-immunoprecipitation

Co-immunoprecipitation was performed using Protein A magnetic beads under non-denaturing conditions. Briefly, A673 and PDX 1460 cells were treated with either vehicle or CX-4945 (2.5 and 5 µM for A673; 1.1 and 2.2 µM for PDX 1460) for 24 h. Cell lysates (90 µg total protein) were precleared with 20 µL of washed Protein A magnetic beads (Cat #73778; Cell Signaling) for 20 minutes at 4°C. Precleared lysates were incubated overnight at 4°C with 5 µg of primary antibody and make up the volume to 200 µL. Species-matched IgG (Cat # 3082; Cell Signaling) served as a negative control. The following day, 25–50 µL of prewashed Protein A magnetic beads were added, and samples were incubated for 20 min at room temperature with rotation. Beads were collected using a magnetic rack and washed 5 times with ice-cold cell lysis buffer. Bound immune complexes were eluted by boiling in 3× Laemmli sample buffer for 5 minutes. An aliquot of 10 µL eluted proteins were resolved by SDS-PAGE and analyzed by immunoblotting with antibodies against the indicated targets.

### Immunoblotting

The ES cell lines and PDX cells were seeded at a density of 4x10^5^ cells/mL in a 6-well plate and left untreated or treated with vehicle (DMSO), CX-4945 at indicated doses for 24 h. The cells were collected by centrifugation, washed with ice-cold PBS and then whole-cell lysates prepared using 1X cell lysis buffer (#9803; Cell Signaling, Danvers, MA) supplemented with 1 mM of PMSF protease Inhibitor (#36978; Thermo Scientific) and phosphatase inhibitor cocktail (#P5726; Sigma-Aldrich). The protein concentration was normalized after performing a bicinchonic acid assay (BCA, Thermo Scientific). After quantification of protein concentration using BCA assay kit (#23225; Thermo Fisher Scientific), an equal amount of protein (10-20 μg) from each sample was denatured at 95°C for 5 min in reducing SDS loading buffer (#7722; Cell Signaling), subjected to gel electrophoresis using 4–20% mini-PROTEAN TGX precast protein gels (Bio-Rad Laboratories, Hercules, CA, RRID:SCR_008426) and transferred to Immobilon-FL PVDF membrane (#IPFL00010; Millipore Sigma). Following blocking with 5% non-fat dry milk in TBST (Tris-buffered saline with 0.1% Tween-20), membranes were incubated with primary antibodies (listed in **Table S5**) overnight at 4°C. After three TBST washes of each 5 min, the blots were incubated with secondary antibodies either anti-rabbit (#7074, Cell Signaling, RRID:AB_2099233) or anti-mouse IgG–horseradish peroxidase (HRP) conjugates (#7076, Cell Signaling, RRID:AB_330924) for 45-60 min at room temperature and washed three times in TBST. SuperSignal West Femto Maximum Sensitivity Substrate (#34095, Thermo Scientific) was used as enhanced chemiluminescent (ECL) substrate and the signals were detected using Azure 500 imaging station (Azure Biosystems, Dublin, CA) or Bio-Rad ChemiDoc MP imaging system (RRID:SCR_008426, Bio-Rad Laboratories, Hercules, CA).

### Organoids Development

Organoids were developed as per described method (60), using the VitroGel ORGANOID (TheWell Bioscience Inc. NJ, USA), that is ready-to-use, xeno-free (animal origin-free) hydrogel system for organoid culture. For ES cell line and ES PDX cell lines 10,000 cells/well were mixed in Organoid formation media in 50 µL volume was overlaid on the solid organoid hydrogels and kept in Co2 Incubator at 37°C for 4-7 days. Using Light microscope (Olympus CKX53 Inverted Microscope (RRID:SCR_025025), individual organoids formed were observed for every 2 days. After 3 days of organoid formation, media was replaced with 50 µL of Organoid expansion media for every 3 days until multiple organoids formed reached the 500-650 µM size. The size of the organoids was measured using Image J software (RRID:SCR_003070).

Briefly, each well in 6 well Glass Chamber slides were coated with 300 µL volume of VitroGel® ORGANOID hydrogels 3. For drug treatment 2X or 4X concentration of drug was added in 150 µL of organoid expansion media. Live/Dead cells in the drug treated organoid were stained using Cyto3D Live-Dead Assay Kit (TheWell Bioscience Inc. NJ, USA) and viewed in confocal microscope (Leica SP8 LIGHTNING confocal microscope (RRID:SCR_018169). 10 µL of Cyto 3D dye was added after 48h of drug treatment and incubated for 30 mins before imaging. Using 63X Water objective lens, individual organoid was imaged in light filter specific to Acridine Orange and Propidium Iodide.

### Lentiviral shRNA transduction

Lentiviral particles containing unique shRNAs for human *CSNK2A1* (#TL320317V) and CK2α overexpression vector (#RC208789L2V) were procured from OriGene Technologies (Rockville, MD, USA). The ES cell lines (A673) were transduced in a 12-well plate at 1-3 MOI (Multiplicity of Infection) using 8 µg/mL of polybrene (#TR-1003-G; Millipore Sigma, Rockville, MD) and spinoculation at 1600 rpm for 60 min at 32°C. The cells were sorted for GFP by flow cytometry under BSL-2 conditions using a BD FACSAria SORP (BD Biosciences, San Jose, CA) instrument in Penn State College of Medicine’s Flow Cytometry Core (RRID:SCR_021134) after 72 h post-transduction and screened for target gene knockdown or overexpression by both qRT-PCR and western blotting. The shRNA with highest knockdown efficiency was selected to achieve high knockdown of *CSNK2A1* gene in the target ES cell lines. The cells transduced with either scramble (Scr) or empty vector (EV) were used as a control for follow-up experiments.

### RNA sequencing analysis

Total RNA was isolated from ES cells (A673, PDX1460) following CX-4945 treatment or CK2α knockdown and paired-end RNA sequencing was performed on the NovaSeq X Plus platform (Novogene, Sacramento, CA). The reads were processed, aligned to the human reference genome (hg38), and visualized as previously described (61). Gene expression levels were normalized and expressed as fragments per kilobase of transcript per million mapped reads (FPKM) using Cufflinks (RRID:SCR_014597) (62). Briefly, differential expression analysis was conducted using edgeR (RRID:SCR_012802), where a quasi-likelihood negative binomial generalized log-linear model was fitted to the data using the glmQLFTest function. Prior to modeling, the raw count data were normalized using the trimmed mean of M-values (TMMs) method. The Benjamini–Hochberg (BH) procedure was applied for multiple testing correction, and genes with an adjusted p-value below 0.05 and fold change above 1.5 were considered as significantly differentially expressed. Hierarchical gene clustering (k-means) analysis with normalized count data and functional enrichment analysis (MSigDB hallmark gene, KEGG) was performed using a web-based application iDEP (http://ge-lab.org/idep/; RRID:SCR_027373) (63). Enrichment of gene signatures for diseases/drugs in differentially expressed genes was analyzed using Enrichr web server (RRID:SCR_001575) (64). The principal component analysis (PCA) and overrepresentation plots were generated using SRplot web platform (RRID:SCR_025904) (65). Gene cluster analysis and heatmaps were visualized using GraphBio (http://www.graphbio1.com/en/) web application (66).

### Flow cytometry

To quantify surface expression, ES cells (A673, SK-ES-1, PDX 1460, PDX 755 and PDX 1287) in culture or lung cells (collected from xenografts) were washed twice with ice-cold PBS and incubated with Zombie Aqua Fixable Viability dye (Biolegend, Cat# 423102) to distinguish Live/Dead cells before primary antibody staining. Cells were washed with FACS staining buffer (Dulbecco’s PBS with 3% FBS) prior to blocking the mouse or human Fc receptor (BD Biosciences, San Jose, CA) for 10 min. Cells were then incubated with respective primary antibodies cocktail [CD99 (BioLegend, Cat# 371313, RRID:AB_2721665), CD133 (BioLegend, Cat# 393906, RRID:AB_2820034) and SEEA4 (BioLegend, Cat# 330406, RRID:AB_1089206)] for 30 minutes on ice in the dark. After washing twice, cells were suspended in FACS buffer and analyzed on a BD LSRFortessa flow cytometer (BD Biosciences, San Jose, CA). The data analysis was performed using the FlowJo v10.10 Software (BD Life Sciences, RRID:SCR_008520). Flow antibodies details are provided in supplemental **Table S6**.

### Animal studies

All mouse care and experiments were conducted in accordance with protocols approved by the Institutional Animal Care and Use Committee (IACUC). A673 and PDX1460 ES cells were injected into 6-8 week-old (15-18g) athymic nude mice (The Jackson Laboratory, Bar Harbor, ME; RRID:IMSR_JAX:002019) on right flank region at a dose of 2500K and 4000K cells per mouse respectively. Cells were suspended in PBS and mixed with the VitroGel Hydrogel Matrix at a 1:1 ratio, and 100uL of mixture was injected per mice. All the healthy weight mice were included in the study, and any mice having illness or weight less than 15g were excluded from the study. Total 20 mice (both sex) randomized into 2 groups, 10 mice per group for each study. Mice were randomized after the one week of cells injection based on the palpable tumor. Group 1 treated with PBS (vehicle) only, Group 2 treated with CX-4945 100 mg/kg via oral gavage twice daily, and treatment continued for the 28-56 days. CX-4945 solution is freshly prepared in sodium phosphate buffer (pH=7.8) and passed through 0.22 µm filter before every dosing. Tumor volume and body weight were measured twice weekly. After the 28 days or when the tumor reached the endpoint, whichever occurred first, mice were sacrificed, and tumor and lung tissues were collected for histology and immunohistochemistry staining. Single cells from both tissues were collected for western blot and flow cytometry analysis. Complete blood count (CBC) and serum clinical chemistry analyses were performed from peripheral blood collected at the time of animal sacrifice.

### Immunohistochemistry (IHC) and H&E analysis

Mouse tumor tissues were obtained, embedded in paraffin, and sectioned at a thickness of 4 μm after being fixed in formalin. After deparaffinization, rehydration, and antigen retrieval, the sections were incubated with a 3% H_2_O_2_ solution for 10 min at room temperature in the dark. After the sections were washed three times with

PBS, they were incubated with 5% goat serum at room temperature for 1 h. Then, the tumor tissue sections were incubated with antibodies working solution prepared in SignalStain antibody diluent (Cell signaling, Cat# 8112) at 4 °C overnight. Details of all antibodies and the dilution used for IHC are described in **Table S7**. The sections were washed and subsequently incubated with a horseradish peroxidase-labelled secondary antibody solution at room temperature for 30 min and visualized using a DAB color development kit. After three minutes of nuclei staining with hematoxylin, images of the histological specimens were captured using a microscope Aperio VERSA ver 8 (RRID:SCR_021016). Hematoxylin and Eosin (H&E) analysis was done. Mitotic figures were counted as a total number identified over 10 consecutive high-power (400x) microscope fields. Counting started at the area with the most mitotic figures. Variance in the mitotic figures may be a result of tumor cell health, blood supply, mutations, effect of treatment, etc. The number may also be artifactually lowered in areas with very high necrosis. Degree of necrosis was quantified as a percentage of the tissue as well as given an overall severity score to assist interpretation. The amount of fibrovascular stroma was also evaluated; the degree of fibrovascular tissue may indicate the ability of the tumor to induce vascularization.

### Statistical analysis

Statistical analyses were performed using GraphPad Prism 10 software (La Jolla, CA; RRID:SCR_002798). For Kaplan-Meier analysis, a log-rank tests were conducted to compare the survival times of treatment group (n=10) versus vehicle group (n=10). Comparisons between two groups were performed using unpaired student’s t-test, whereas comparisons among three or more groups were tested by analysis of variance (ANOVA) with suitable post hoc tests as indicated in each figure legend. All data are represented as the mean ± standard error of the mean (SEM) or standard deviation (SD). A value of p<0.05 was considered statistically significant. The significance level of p-values is indicated by asterisks (*P<0.05; **P<0.01; ***P<0.001; ****P<0.0001; ns, not significant).

## Authors’ Contributions

**M. Daniyal:** Resources, data curation, formal analysis, validation, investigation, visualization, methodology, writing–original draft, writing–review and editing. **R. Rajaiah:** Resources, data curation, formal analysis, validation, investigation, visualization, methodology, writing–original draft, writing–review and editing. **U. Golla:** Data curation, formal analysis, investigation, visualization, methodology, writing–original draft, writing–review and editing. **M. Shanmugam:** Resources, formal analysis, validation, investigation, methodology, writing–review and editing. **R. C. Sholler**, **J. Hengst**, and **A.B. Nagulapally:** Resources, formal analysis, writing–review and editing. **H. Valensi:** Software, formal analysis, visualization, writing–review and editing. **L. Matthew:** Formal analysis, visualization, writing–review and editing. **Y. Uzun:** Data curation, software, formal analysis, visualization, writing–review and editing. **G.S. Sholler:** Formal analysis, writing–review and editing. **C.G. Behura:** Conceptualization, resources, data curation, formal analysis, supervision, funding acquisition, writing–original draft, project administration, writing–review and editing.

## Acknowledgments

The authors would like to thank the Beat Childhood Cancer Consortium and the biorepository for sharing the Ewing sarcoma patient-derived xenograft (PDX) cells. We used Grammarly’s AI features to proofread this document for spelling and grammar errors and to revise text to improve clarity and readability.

## Institutional Review Board Statement

All the animal experiments were conducted in accordance with guidelines and following protocols approved by the Institutional Animal Care and Use Committee at Penn State Hershey, Hershey, PA (IACUC # PROTO201901034). De-identified patient-derived xenograft (PDX) samples were provided by collaborators at the Penn State Cancer Institute and the Beat Childhood Cancer biorepository at Penn State University College of Medicine (Hershey, PA, USA), and used in compliance with Institutional Review Board regulations.

## Informed Consent Statement

Not applicable.

