## Supplementary material for "CK2 inhibitor CX-4945 targets EWS-FLI1 signaling network and shows therapeutic efficacy in metastatic mouse models of Ewing Sarcoma": Figure S1-S6 and Table S1-S7

**Running title:** CK2, a novel therapeutic target in Ewing sarcoma

Muhammad Daniyal[1,†], Rajesh Rajaiah[1a,†], Upendarrao Golla[1], Marudhu Pandiyan Shanmugam[1], Chloe Sholler[2], Jeremy Hengst[1,3], Abhinav B. Nagulapally[2], Hannah Valensi[1], Lanza Matthew[4], Yasin Uzun[1,3], Giselle Saulnier Sholler[1,3], Chandrika Gowda Behura[1*]

[1]Department of Pediatrics, Pennsylvania State University College of Medicine, Hershey, PA USA

[2]Penn State Health Children's Hospital and Penn State College of Medicine, Hershey, PA USA

[3]Department of Molecular and Precision Medicine, Pennsylvania State University College of Medicine, Hershey, PA USA

[4]Department of Comparative Medicine, Pennsylvania State University College of Medicine, Hershey, PA USA

[a]Present address: NorthEast BioLab, Hamden, CT 06518.

[†]These authors contributed equally

***Corresponding Author:** Dr. Chandrika Gowda Behura

### Supplementary Figures

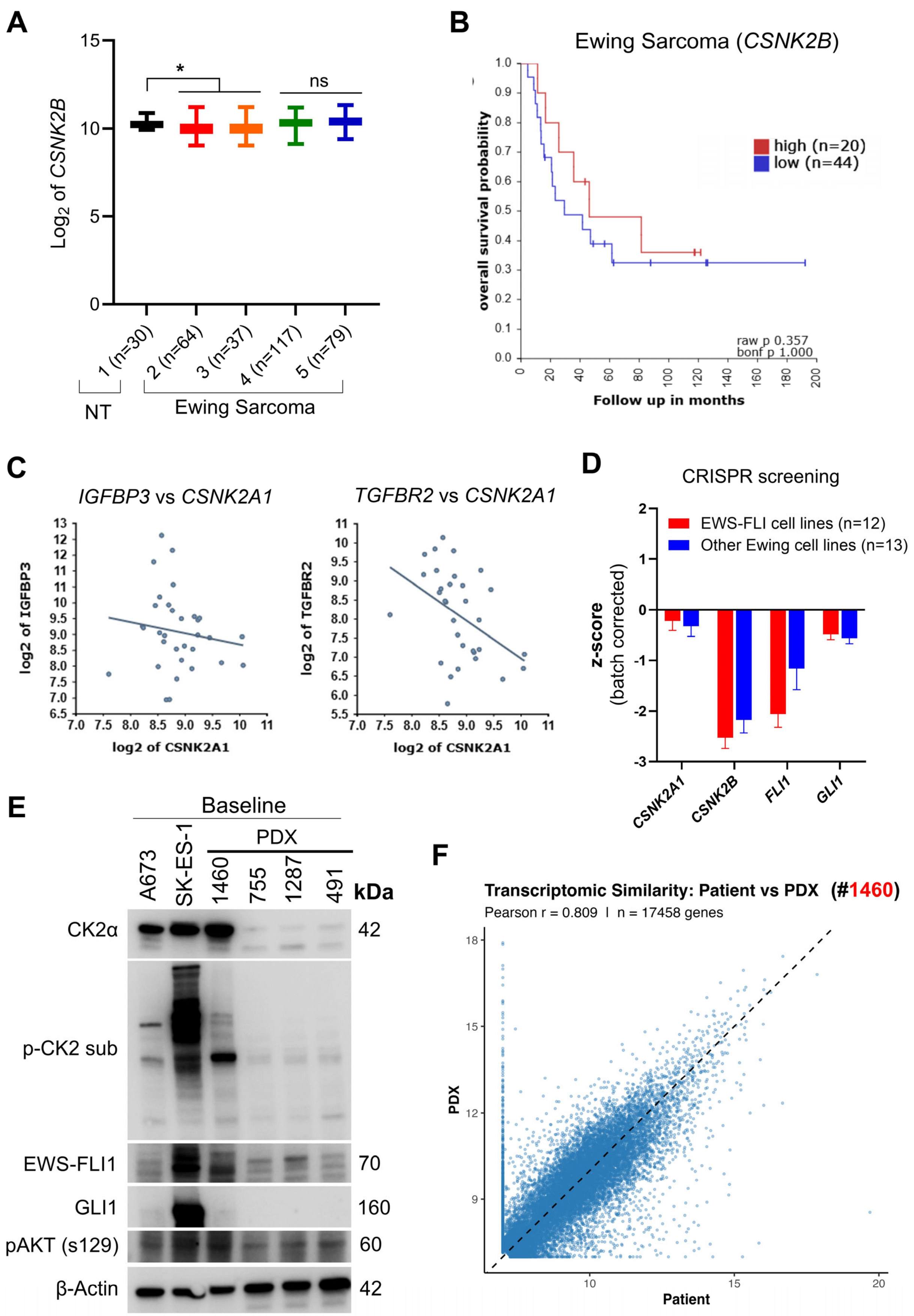

**Supplementary Figure 1: Correlation between *CSNK2A1* and other Ewing sarcoma specific genes. A)** *CSNK2B* mRNA expression in primary Ewing sarcoma (ES) patient samples (aggregated from four datasets in R2 platform and listed in **Table S1**) compared to normal tissue (NT; n=30). *$p<0.05$ by mixed-effects analysis (Dunnett's multiple comparisons test) indicates significant difference compared to NT group. **B)** Kaplan–Meier survival analysis of ES patients stratified by high versus low *CSNK2B* expression. Data were analyzed using the R2: Genomics Analysis and Visualization Platform (https://r2.amc.nl/), with groups defined by median expression cutoff (n=64; GSE17679 accessed from R2 platform). Statistical significance was determined by the log-rank test ("bonf p" refers to the Bonferroni-corrected p-value). **C)** Correlation between *CSNK2A1* (CK2α), *IGFBP3* and *TGFBR2* gene expression levels from Ewing sarcoma patient samples (n=64; GSE17679 accessed from R2 platform). **D)** CRISPR screen scores (batch corrected) for indicated genes of interest in Ewing sarcoma cell lines without (n=13) and with EWS-FLI1 fusion (n=12). Negative z-scores indicate that targeting a specific gene caused a decrease in cell viability (dropout) and is essential in Ewing sarcoma cells. *FLI1* and *GLI1* genes that are crucial for Ewing sarcoma cells survival were used as controls. The CRISPR screening data for Ewing sarcoma cells was obtained from iCSDB database (https://www.kobic.re.kr/icsdb/). **E)** Immunoblotting analysis of baseline protein expression for CK2α, EWS-FLI1, and GLI1, alongside CK2 catalytic activity (p-AKT Ser129), in the indicated ES cell lines and diverse ES patient-derived xenografts (PDX: 1469, 755, 1287, 491). Blots from one of the representative experiments are shown. **F)** Comparison of variance stabilizing transformation (VST) RNA-seq expression values between the patient tumor (PT) and the mean of three vehicle-treated PDX replicates. Pearson correlation (*r*) used to measure the strength and direction of a linear relationship.

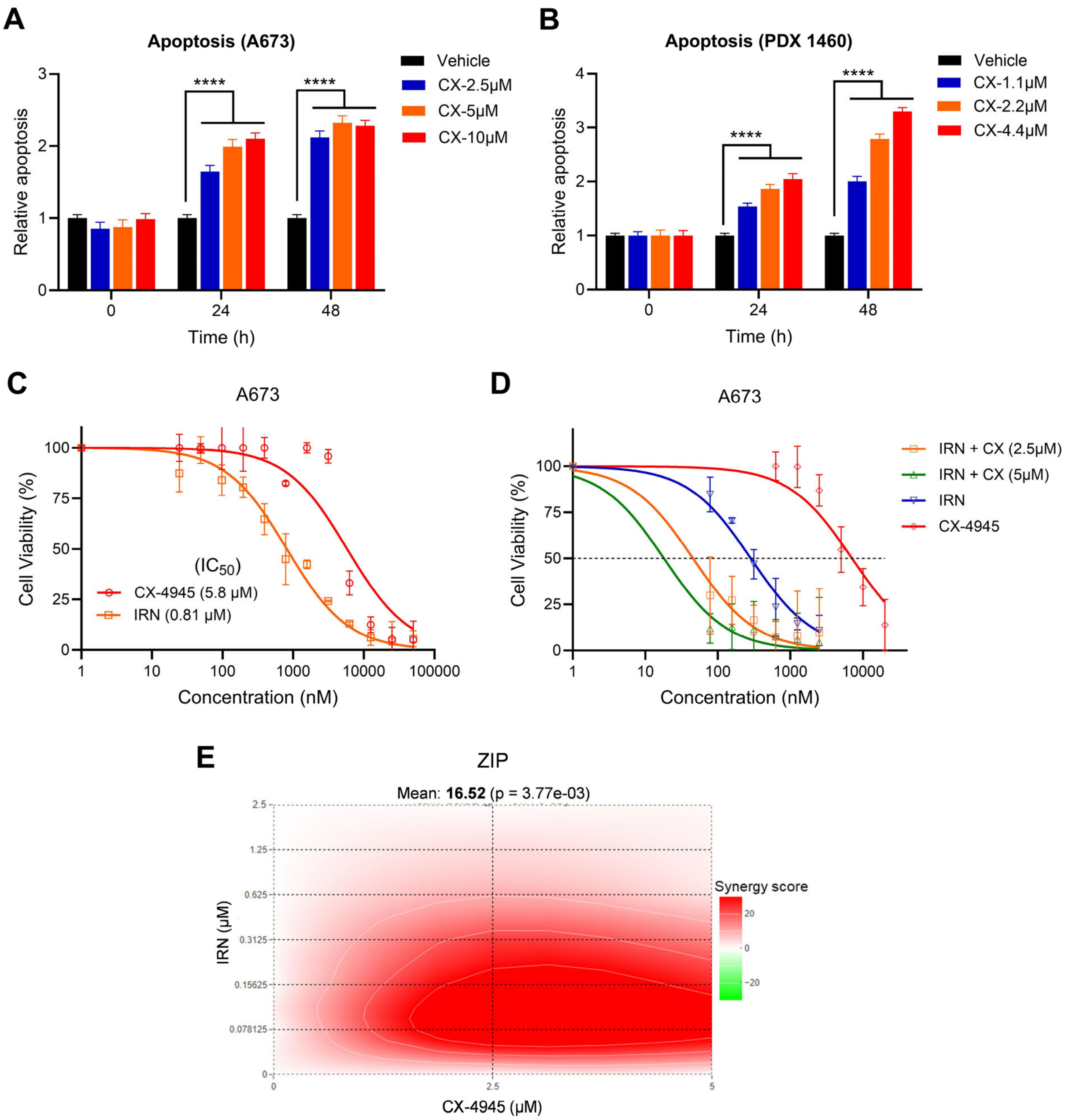


**Supplementary Figure 2: CX-4945 treatment promotes apoptosis and synergizes with Irinotecan in Ewing Sarcoma cells. A-B)** Ewing sarcoma A673 cell line (A) and PDX1460 cells (B) were treated with different concentrations of CX-4945 (CX) for 48 h and assessed for cell apoptosis using Incucyte Caspase 3/7 green dye. The data are presented as mean ± SD (n=6) relative to vehicle (DMSO) control cells and analyzed by two-way ANOVA (Dunnett's multiple comparisons test). ****p<0.0001 denotes statistical significance. **C-D)** A673 Ewing sarcoma cell line was treated with different concentrations of CX-4945 (CX) and Irinotecan (IRN) alone (**C**) or in combination with indicated doses for 48 h (**D**). Cell viability was assessed using WST reagent and cell viability was calculated relative to vehicle-treated cells as 100%. **E)** ZIP synergy heatmap and score for CX-4945 and Irinotecan (IRN) combination in A673 ES cell line was shown. ZIP scores greater than 10 indicates 'synergy', and 0-10 indicate 'additive' activity between CX-4945 and IRN.

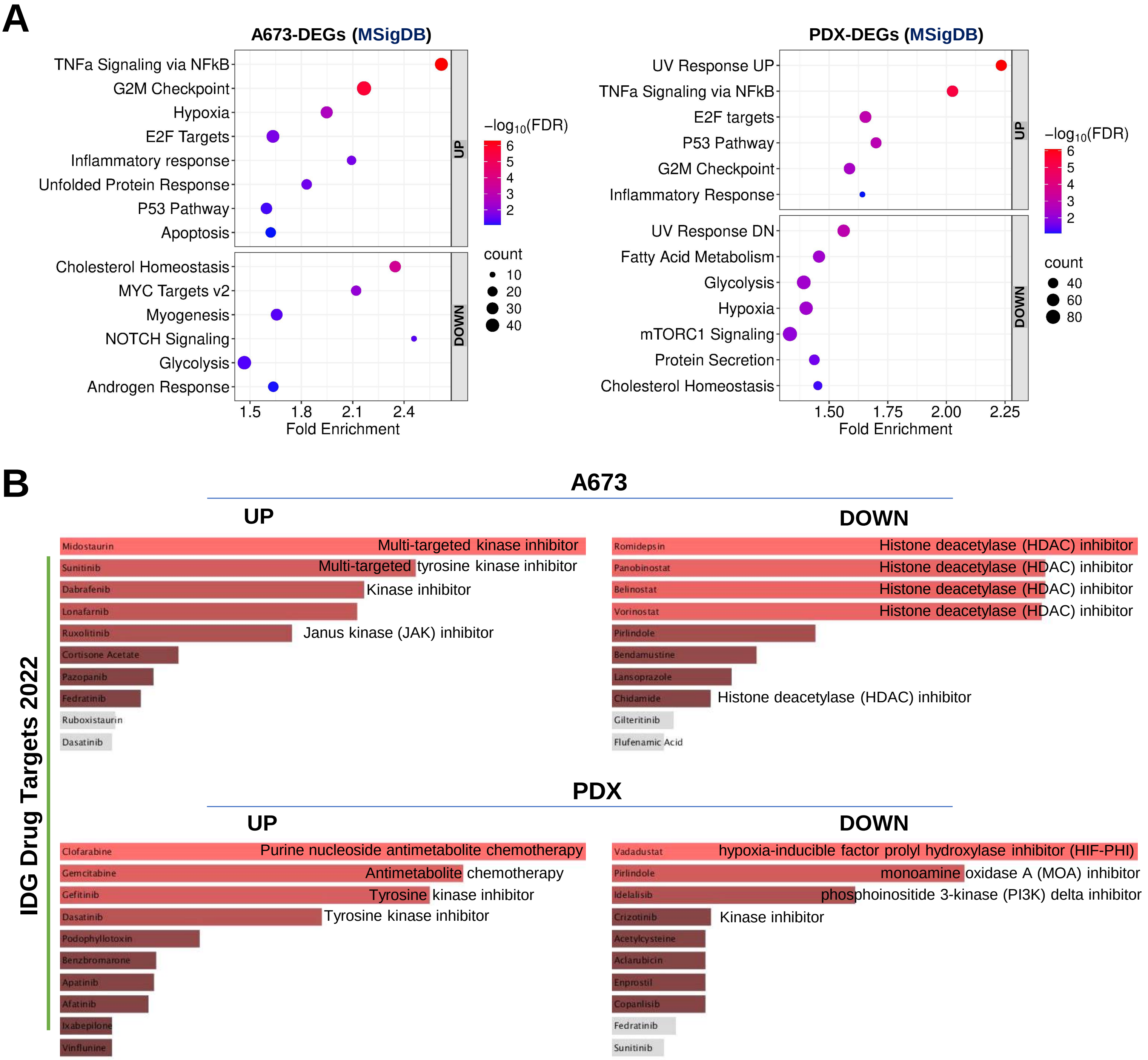


**Supplementary Figure 3: Overrepresentation analysis of Ewing sarcoma cells transcriptome after CX-4945 treatment. A)** Dot plot displaying the top-ranked functional pathways (MSigDB hallmark gene set) overrepresented in differentially expressed genes (UP- and DOWNregulated) following CX-4945 treatment of ES cells (A673, PDX1460). **B)** DrugSeq (Drug perturbation signatures) enrichment analysis bar plot of DEGs obtained by Enrichr web tool. The genes upregulated and downregulated in A673 and PDX1460 Ewing sarcoma cells after 24h treatment with CX-4945 were checked against IDG_DRUG_Targets_2022 gene-set libraries that significantly overlaps with input genes. The length of the bar and intensity of the color represents the significance of that specific gene-set or term.

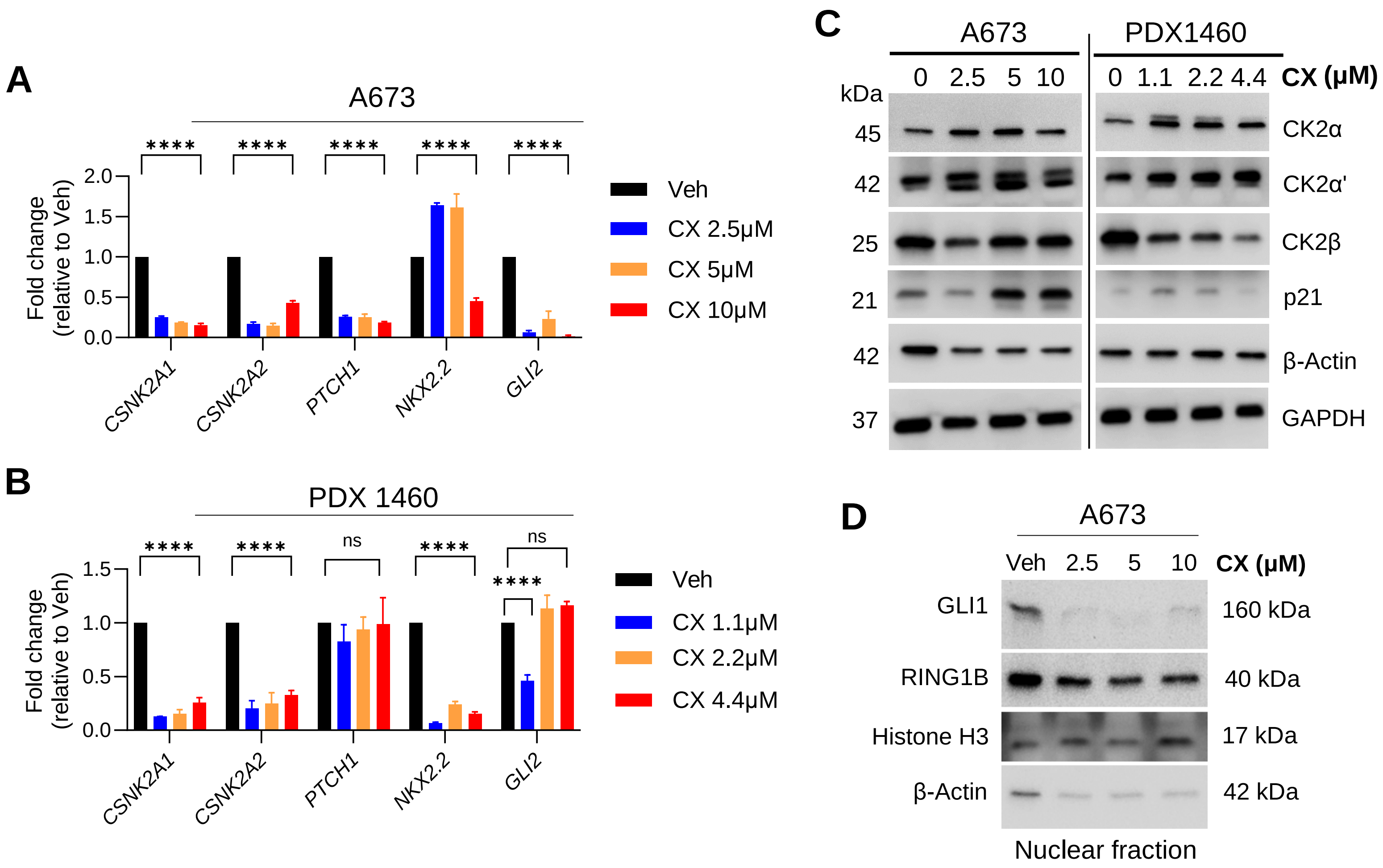


**Supplementary Figure 4: The effect of CX-4945 treatment on the expression of CK2 and EWS-FLI1 target genes. A-B)** A673 (**A**) and PDX-1460 (**B**) were treated with different indicated concentration of CX-4945 and gene expression of CK2 (CSNK2A1, CSNK2A2) and EWS-FLI1 target genes (key mediators) was analyzed by qRT-PCR. The data are presented as mean ± SD (n=3 replicates from a representative run) relative to vehicle (DMSO) control cells. ****p<0.0001 by two-way ANOVA (Dunnett's multiple comparisons test) indicates statistical significance. 'ns' denotes 'not significant'. **C)** Immunoblot analysis of CK2 protein kinase subunits after CX-4945 treatment. A673 cells (left) and PDX 1460 cells (right) were treated with increasing concentrations of CX-4945 for 24 h. Whole-cell lysates were assessed for CK2α, CK2α', CK2β, and p21 levels. **D)** Western blot analysis of GLI1 and RING1B levels in the nuclear fraction isolated from A673 cells after treatment with vehicle (DMSO) and indicated doses of CX-4945 for 24 h. β-Actin (cytoplasmic) and histone H3 (nuclear) antibodies were used to confirm successful cell fractionation. Blots from one of the representative experiments are shown in '**C** & **D**'.

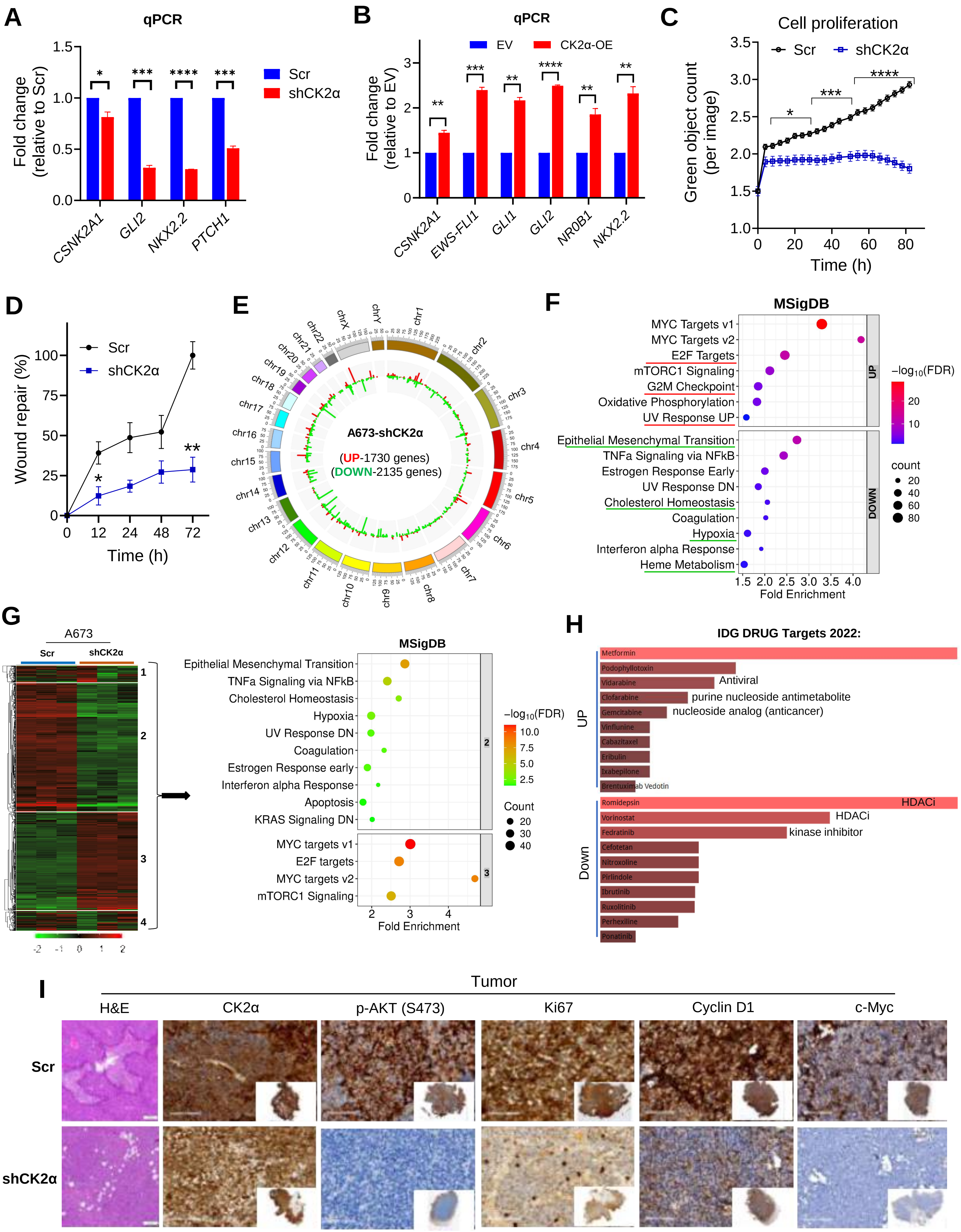
A
qPCR
Fold change
(relative to Scr)
1.5
1.0
0.5
0.0
*
***
****
***
Scr
shCK2α
CSNK2A1
GLI2
NKX2.2
PTCH1
B
qPCR
EV
CK2α-OE
Fold change
(relative to EV)
3
2
1
0
**
***
**
****
**
**
CSNK2A1
EWS-FLI1
GLI1
GLI2
NR0B1
NKX2.2
C
Cell proliferation
Scr
shCK2α
Green object count
(per image)
3.5
3.0
2.5
2.0
1.5
1.0
*
***
****
0
20
40
60
80
Time (h)
D
Wound repair (%)
100
75
50
25
0
Scr
shCK2α
*
**
0
12
24
48
72
Time (h)
E
A673-shCK2α
(UP-1730 genes)
(DOWN-2135 genes)
chr1
chr2
chr3
chr4
chr5
chr6
chr7
chr8
chr9
chr10
chr11
chr12
chr13
chr14
chr15
chr16
chr17
chr18
chr19
chr20
chr21
chr22
chrX
chrY
F
MSigDB
MYC Targets v1
MYC Targets v2
E2F Targets
mTORC1 Signaling
G2M Checkpoint
Oxidative Phosphorylation
UV Response UP
Epithelial Mesenchymal Transition
TNFa Signaling via NFkB
Estrogen Response Early
UV Response DN
Cholesterol Homeostasis
Coagulation
Hypoxia
Interferon alpha Response
Heme Metabolism
UP
DOWN
−log10(FDR)
20
10
count
20
40
60
80
1.5
2.0
2.5
3.0
3.5
4.0
Fold Enrichment
G
A673
Scr
shCK2α
1
2
3
4
-2
-1
0
1
2
MSigDB
Epithelial Mesenchymal Transition
TNFa Signaling via NFkB
Cholesterol Homeostasis
Hypoxia
UV Response DN
Coagulation
Estrogen Response early
Interferon alpha Response
Apoptosis
KRAS Signaling DN
MYC targets v1
E2F targets
MYC targets v2
mTORC1 Signaling
2
3
−log10(FDR)
10.0
7.5
5.0
2.5
Count
20
30
40
2
3
4
Fold Enrichment
H
IDG DRUG Targets 2022:
UP
Metformin
Podophyllotoxin
Vidarabine
Antiviral
Clofarabine
purine nucleoside antimetabolite
Gemcitabine
nucleoside analog (anticancer)
Vinflunine
Cabazitaxel
Eribulin
Ixabepilone
Brentuximab Vedotin
Down
Romidepsin
HDACi
Vorinostat
HDACi
Fedratinib
kinase inhibitor
Cefotetan
Nitroxoline
Pirlindole
Ibrutinib
Ruxolitinib
Perhexiline
Ponatinib
I
Tumor
H&E
CK2α
p-AKT (S473)
Ki67
Cyclin D1
c-Myc
Scr
shCK2α

**Supplementary Figure 5: Genetic knockdown of CK2α inhibits migration and promote apoptosis by alteration of transcriptome in A673 cells. A and B)** Quantification of CK2α (*CSNK2A1*) and EWS-FLI1 target genes after CK2α knockdown (**A**) and overexpression (**B**) in A673 Ewing sarcoma cells by qRT-PCR. The data are presented as mean ± SD (n=3 replicates from a representative run) relative to Scr (scramble) or EV (empty vector) control cells and analyzed by unpaired t-test (Welch's correction). **C)** Green optimal count at different time points was recorded in A673 cells (GFP positive) expressing either scramble (Scr) control or CK2α-specific shRNA (shCK2α) by Incucyte. The data are presented as mean ± SEM (n=6) and analyzed by unpaired t-test (Welch's correction). **D)** Summary graph showing wound healing rates by A673 cells expressing either scramble (Scr) control or CK2α-specific shRNA (shCK2α). The data are presented as mean ± SEM (n=3) and analyzed by unpaired t-test (Welch's correction). *p<0.05, **p<0.01, ***p<0.001, and ****p<0.001 denotes statistical significance in **A-D**. **E**) Circos plot showing the global transcriptomic shift in *CSNK2A1*-knockdown A673 cells; red and green bars indicate significantly upregulated and downregulated genes, respectively (adjusted p-values <0.05; fold change >1.5). **F)** Dot plot displaying the top-ranked functional pathways (MSigDB hallmark gene set) overrepresented in differentially expressed genes (UP- and DOWNregulated) following CK2α knockdown in A673 ES cells. **G)** Heatmap showing the hierarchical clustering (k-means) of most variable genes (top 2000) in A673 cells transcriptome after CK2α knockdown. Subsequent overrepresentation analysis of top-ranked functional pathways (MSigDB hallmark gene set) in different gene clusters obtained with k-means clustering were showed in dot plot. **H)** DrugSeq (Drug perturbation signatures) enrichment analysis bar plot of DEGs obtained by Enrichr web tool. The genes upregulated and downregulated after CK2α knockdown in A673 Ewing sarcoma cells were checked against IDG_DRUG_Targets_2022 gene-set libraries that significantly overlaps with input genes. The length of the bar and intensity of the color represents the significance of that specific gene-set or term. **I**) Histopathological evaluation of A673 xenograft tumors by H&E staining and immunohistochemistry (IHC). Representative images of primary tumor sections from Scr and shCK2α groups stained for CK2α, p-AKT (S473), Ki67 (proliferation marker), Cyclin D1, and c-Myc. Inset image shows entire view of the tissue section.

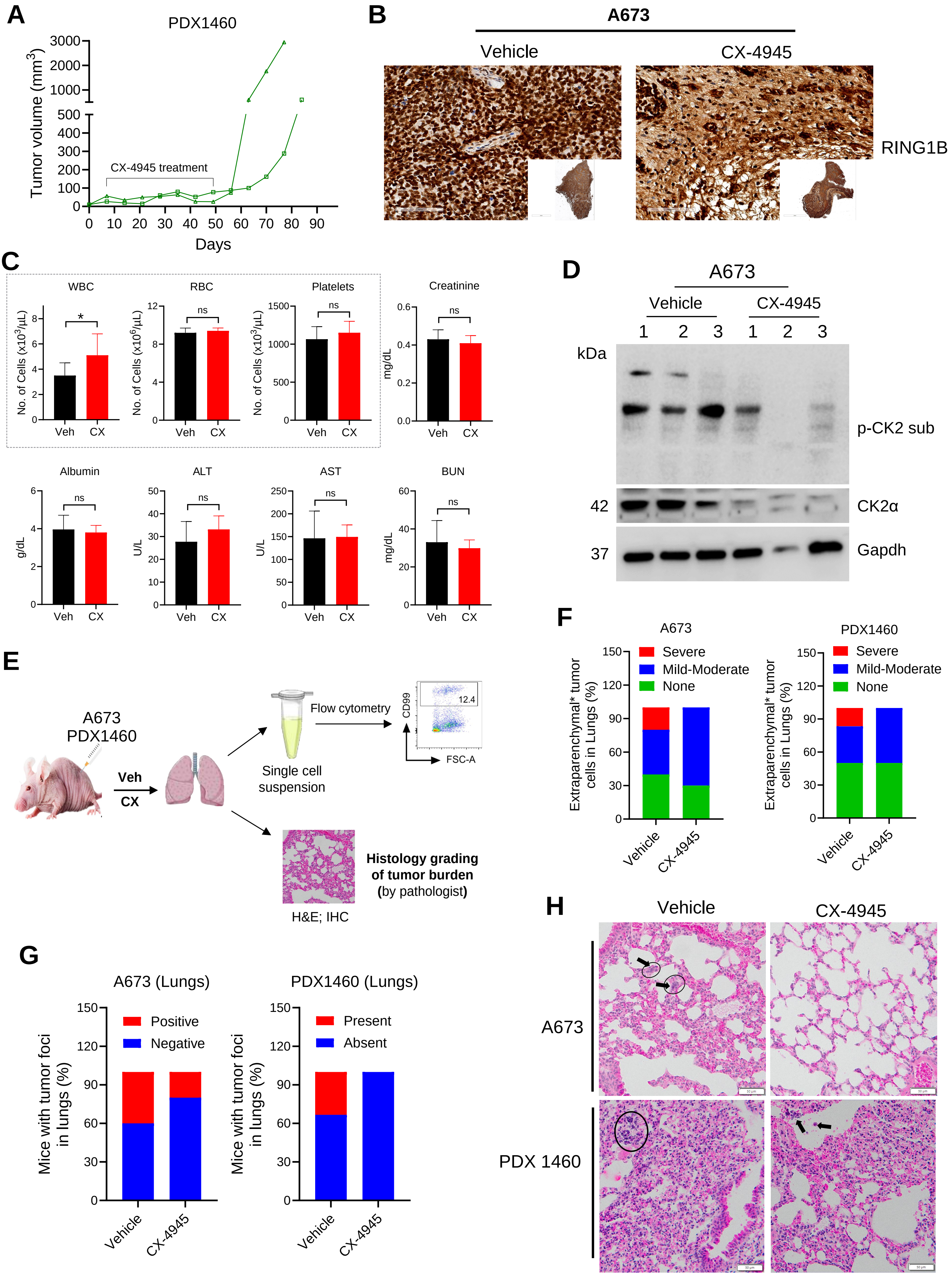
A
PDX1460
Tumor volume (mm3)
CX-4945 treatment
Days
B
A673
Vehicle
CX-4945
RING1B
C
WBC
RBC
Platelets
Creatinine
No. of Cells (x10^3/µL)
No. of Cells (x10^6/µL)
mg/dL
Veh
CX
ns
Albumin
ALT
AST
BUN
g/dL
U/L
D
A673
Vehicle
CX-4945
kDa
p-CK2 sub
42
CK2α
37
Gapdh
E
A673
PDX1460
Veh
CX
Flow cytometry
CD99
12.4
FSC-A
Single cell suspension
Histology grading of tumor burden (by pathologist)
H&E; IHC
F
A673
PDX1460
Severe
Mild-Moderate
None
Extraparenchymal* tumor cells in Lungs (%)
Vehicle
CX-4945
G
A673 (Lungs)
PDX1460 (Lungs)
Positive
Negative
Present
Absent
Mice with tumor foci in lungs (%)
H
Vehicle
CX-4945
A673
PDX 1460

**Supplementary Figure 6: CX-4945 inhibits lung metastasis in Ewing sarcoma *in vivo* mouse models. A)** Longitudinal monitoring of tumor volume in PDX1460 xenografts (pilot study with n=2), showing a significant reduction in tumor burden following 6-weeks of CX-4945 treatment. Discontinuation of CX-4945 treatment resulted in rapid tumor growth. **B)** The tumors collected from A673 xenograft mice after vehicle and CX-4945 treatment at study termination were processed for immunohistochemistry (IHC). Representative IHC images of primary tumor sections stained with RING1B are presented. Inset image shows entire view of the tissue section. **C)** CBC and serum clinical chemistry analysis of A673 mice after 3-4 weeks of treatment with either vehicle or CX-4945. The data are presented as mean ± SD (n=10). *$p<0.05$ by unpaired t-test (Welch's correction) indicates statistical significance and 'ns' denotes 'not significant'. **D)** Immunoblotting analysis of CK2α and its substrate phosphorylation in tumors harvested from A673 xenograft mice (vehicle or CX-4945) at study termination. Gapdh was used as a loading control. Tumors collected from three different mice from each cohort are processed and presented. **E)** Schematic diagram illustrating the procedure for collection and processing of lung tissue from A673 or PDX1460 xenografts after treatment with vehicle (Veh) or CX-4945 (CX) for flow cytometric analysis or histology grading of tumor burden by a pathologist. **F and G)** Lung tissues collected at the study endpoint were fixed and embedded in paraffin blocks. Lung tissue section slides prepared from paraffin blocks were analyzed for tumor burden by a pathologist after H&E staining. The pathologist was blinded to the group assignment during review. At least 2-3 slides from each lung tissue were reviewed. The number of small, round, blue cell tumor foci (organized cluster of tumor cells) in lung tissue was counted, and the size was measured. The extra-parenchymal tumor cells were semi-quantitatively graded as none, mild, moderate or severe by the pathologist. The data are presented as percentage of mice from each study cohort (vehicle or CX-4945; n=10 for A673 and n=6 for PDX1460) showed different grades of extra-parenchymal tumor cells in the lungs (**F**). The percentage of mice in each group that contain tumor foci in lungs are shown in (**G**). **H)** Representative H&E stained images of lung tissue (bearing tumor foci) from vehicle and CX-4945 treated ES xenografts.

**Supplementary Tables**

**Table S1:** List of datasets used from R2 platform*

| S.No. | Cells | Database | Number of Samples | GEO ID Accession number | Pubmed ID |
|---|---|---|---|---|---|
| 1 | Normal Bonemarrow Mesenchymal stem cells | Yamaguchi | 30 | GSE7637 | 18550633 |
| 2 | Ewing Sarcoma | Savola | 64 | GSE17679 | 22084725 |
| 3 | Ewing Sarcoma | Francesconi | 37 | GSE12102 | 19307502 |
| 4 | Ewing Sarcoma | Delattre | 117 | GSE34620 | 22327514 |
| 5 | Ewing Sarcoma | Surdez | 79 | GSE142162 | 33930311 |

*These studies were used to generate **Figure 1A-E** and **Figure S1A-C**.

**Table S2:** Characteristics of Ewing Sarcoma cell lines

| Cell line | Source | Cat. No. | Tissue | Age | Gender | Gene mutation/ Fusion |
|---|---|---|---|---|---|---|
| A673 | ATCC | CRL-1598 | Muscle | 15 y | F | EWS-FLI1, BRAF, TP53 |
| SK-ES-1 | ATCC | HTB-86 | Bone | 18 y | M | EWS-FLI1, STAG2, TP53 |

**Table S3:** Characteristics of Ewing sarcoma patient-derived xenograft (PDX) samples

| S.No. | PDX ID | Age | Gender | EWS Fusion | *CSNK2A1* (TPM) | Tumor /biopsy site (or) BM aspirate |
|---|---|---|---|---|---|---|
| 1 | 1460 | 13 y | F | EWS-FLI | 6.21 | Right Thoracic mass |
| 2 | 755 | 12 y | F | EWS-FLI | 6.18 | Lung |
| 3 | 1287 | 18 y | M | Wild-type | 6.43 | Metastasis |
| 4 | 491 | 14 y | F | EWS-FLI | 5.63 | Bone/Femur |
| 5 | 65 | 15 y | M | EWS-FLI | 6.38 | N/A |
| 6 | 1258 | 21 y | F | EWS-FLI | 4.44 | Metastasis |
| 7 | 1251 | 11 y | M | EWS-ERG | 5.27 | Rib |
| 8 | 1306 | 19 y | M | EWS-FLI | 4.17 | Left proximal humerus |
| 9 | 62 | 11 y | M | EWS-FLI | 6.11 | N/A |
| 10 | 198 | 13 y | F | N/A | N/A | N/A |
| 11 | 634 | 18 y | F | EWS-FLI | 6.42 | Retroperitoneum |
| 12 | 58 | 13 y | F | N/A | N/A | Left Ilium |
| 13 | 1050 | 18 y | F | Wild-type | 2.8 | N/A |
| 14 | 416 | 8 y | M | EWS-FLI | 5.92 | Bone/Femur |
| 15 | 1419 | 15 y | F | EWS-FLI | 7.46 | Pelvic mass |

Ewing sarcoma PDX cells used for immunophenotyping and immunoblotting analysis are highlighted using red-colored box. 'N/A' denotes 'not available'.

**Table S4:** List of primers used for qRT-PCR

| Target | Forward primer (5' → 3') | Reverse primer (5' → 3') | Source |
|---|---|---|---|
| *CSNK2A1* | GGTGAGGATAGCCAAGGTTCTG | TCACTGTGGACAAAGCGTTCCC | IDT |
| *CSNK2A2* | CGACCATCAACAGAGACTGACTG | GTGAGACCACTGGAAAGCACAG | IDT |
| *EWS–FLI1* | CAGTCACTGCACCTCCATCC | TTCATGTTATTGCCCCAAGC | IDT |
| *GLI1* | GAACCTTCCTACCAGAGTCC | GTGCTGCTGCCCTATGTG | IDT |
| *GLI2* | AGATGTTGTAAGAGAAGGTTTATG | CGTTAGCCGAATGTCAGC | IDT |
| *PTCH1* | ACAAACTCCTGGTGCAAACC | CTTTGTCGTGGACCCATTCT | IDT |
| *NKX2.2* | CTACGACAGCAGCGACAACC | GCCTTGGAGAAAAGCACTCG | IDT |
| *RING1B* | CAGACAAACGGAACTCAACCATT | CTGTTATTGCCTCCTGAGGTGTT | IDT |
| *EZH2* | AAGAAATCTGAGAAGGGACC | CTCTTTACTTCATCAGCTCG | IDT |
| *NR0B1* | CAGTCAGCATGGATGATATG | CACAGCTCTTTATTCTTCCC | IDT |
| *IGF1* | CCCAGAAGGAAGTACATTTG | GTTTAACAGGTAACTCGTGC | IDT |
| *IGFBP3* | CTGCTCAGATTTCCCCAAAG | TGGCATCAAGCAGGTCATAG | IDT |
| *TGFBR2* | CATCTGTGAGAAGCCACAGG | TGCACTCATCAGAGCTACAGG | IDT |
| *GAPDH* | GTCTCCTCTGACTTCAACAGCG | ACCACCCTGTTGCTGTAGCCAA | IDT |

**Table S5:** List of antibodies used for immunoblotting analysis of Ewing sarcoma cells

| Target | Clone | Reference | Provider |
|---|---|---|---|
| CK2α | E-7 | SC-373894 | Santa Cruz Biotechnology |
| CK2α' | D-7 | SC-514403 | Santa Cruz Biotechnology |
| CK2β | EPR1994 | Ab133576 | Abcam |
| p-CK2 sub | | 8738s | Cell Signaling |
| p-AKT (S129) | D4P7F | 13461s | Cell Signaling |
| p-AKT (S473) | 193H12 | 4058s | Cell Signaling |
| Akt (pan) | | 9272s | Cell Signaling |
| β-Catenin | 6B3 | 9582s | Cell Signaling |
| Cyclin D1 | E3P5S | 55506s | Cell Signaling |
| c-Myc | E5Q6W | 18583S | Cell Signaling |
| p21 | 12D1 | 2947s | Cell Signaling |
| GLI1 | C68H3 | 3538s | Cell Signaling |
| FLI1 (to detect EWS-FLI1) | | Ab180902 | Abcam |
| EWSR1 (to detect EWS-FLI1) | | A300-417 | Bethyl Laboratories |
| RING1B | | D139-3 | Medical & Biological Laboratories |
| PARP | | 9542s | Cell Signaling |
| Histone H3 | | 9715s | Cell Signaling |
| Ubiquitin | E4I2J | 43124s | Cell signaling |
| K48-linkage Specific Polyubiquitin | D9D5 | 8081s | Cell signaling |
| β-actin | 13ES | 4970s | Cell Signaling |
| Gapdh | 14C10 | 2118 | Cell Signaling |

**Table S6:** List of antibodies used for flow cytometry analysis of Ewing sarcoma cells

| Target | Fluorochrome | Clone | Reference | Species | Provider |
|---|---|---|---|---|---|
| hCD45 | APC/Cy7 | HI30 | 304014 | Mouse IgG1, κ | BioLegend |
| hCD99 | PE/Cy7 | 3B2/TA8 | 371313 | Mouse IgG2a κ | BioLegend |
| hCD133 | APC | S16015F | 393906 | Mouse IgG2a, κ | BioLegend |
| hSSEA-4 | PE | MC-813-70 | 330406 | Mouse IgG3, κ | BioLegend |

**Table S7:** List of antibodies used for Immunohistochemistry (IHC) assay

| Target | Clone | Dilution | Reference | Provider |
|---|---|---|---|---|
| Ki67 | K2 | Pre-diluted | PA0230-U | Leica Biosystems |
| CD99 | PCB1 | 1:50 | CD99-187-L-U | Leica Biosystems |
| CK2 | | 1:1600 | N/A | Senhwa Biosciences |
| RING1B | 3-3 | 1:200 | D139-3 | Medical & Biological Laboratories |
| FLI1 (to detect EWS-FLI1) | EPR4646 | 1:100 | Ab133485 | Abcam |
| Akt (pan) | C67E7 | 1:600 | 4691s | Cell Signaling |
| p-AKT (S129) | D4P7F | 1:200 | 13461s | Cell Signaling |
| p-AKT (S473) | D9E | 1:200 | 4060s | Cell Signaling |
| β-Catenin | 6B3 | 1:200 | 9582s | Cell Signaling |
| Cyclin D1 | E3P5S | 1:200 | 55506s | Cell Signaling |
| c-Myc | E5Q6W | 1:200 | 18583s | Cell Signaling |
